# RET Signaling Persists in the Adult Intestine and Stimulates Motility by Limiting PYY Release from Enteroendocrine Cells

**DOI:** 10.1101/2022.04.15.488514

**Authors:** Amy Shepherd, Laurence Feinstein, Svetlana Sabel, Daniella Rastelli, Esther Mezhibovsky, Lynley Matthews, Anoohya Muppirala, Ariel Robinson, Michael D. Gershon, Meenakshi Rao

## Abstract

**Background & Aims:** RET receptor tyrosine kinase is necessary for enteric nervous system (ENS) development. Loss-of-function *RET* mutations cause Hirschsprung disease (HSCR), in which infants are born with aganglionic bowel. Despite surgical correction, HSCR patients often experience chronic defecatory dysfunction and enterocolitis, suggesting that RET is important after development. To test this hypothesis, we determined the location of postnatal RET and its significance in gastrointestinal (GI) motility.

**Methods:** *Ret*^CFP/+^ mice and human transcriptional profiling data were studied to identify the enteric neuronal and epithelial cells that express RET. To determine whether RET signaling in these cells regulates adult gut motility *in vivo*, genetic and pharmacologic approaches were used to disrupt RET in either all RET-expressing cells, a major subset of enteric neurons, or intestinal epithelial cells.

**Results:** Distinct subsets of enteric neurons and enteroendocrine cells expressed RET in the adult intestine. RET disruption in the intestinal epithelium, rather than in enteric neurons, slowed GI motility selectively in adult male mice. This effect was phenocopied by RET kinase inhibition. Most RET^+^ epithelial cells were either enterochromaffin cells that release serotonin (5-HT) or L-cells that release peptide YY (PYY), both of which can alter motility. RET kinase inhibition exaggerated PYY release in a nutrient-dependent manner without altering 5-HT secretion. PYY receptor blockade fully rescued dysmotility in mice lacking epithelial RET.

**Conclusion:** RET signaling normally limits nutrient-dependent PYY release from L-cells and this activity is necessary for normal intestinal motility in male mice. These effects could contribute to post-operative dysmotility in HSCR, which predominantly affects males, and uncovers a mechanism that could be targeted to treat post-prandial GI dysfunction.

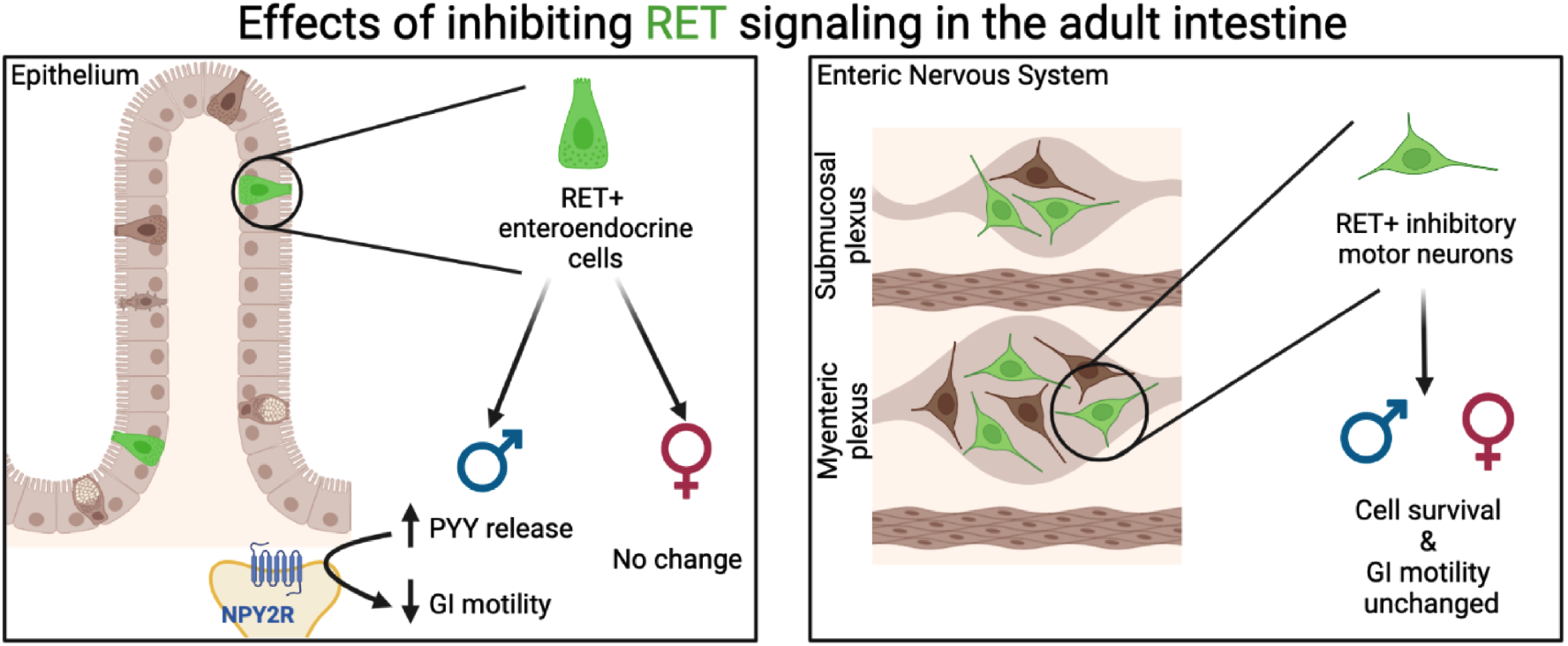

## Introduction

RET tyrosine kinase is a transmembrane protein that functions as the signaling co-receptor for several secreted ligands. RET is a proto-oncogene and activating mutations are linked to neoplasia. In contrast, inactivating RET mutations cause developmental defects, most notably Hirschsprung’s disease (HSCR). HSCR is a male-predominant disorder in which infants are born lacking enteric ganglia, typically in the distal colon. RET dysfunction is a major cause of both familial and sporadic forms of HSCR with more than 60% of patients found to have pathogenic alleles leading to diminished RET signaling^1^. The neural crest derived progenitors that colonize the bowel require RET for normal migration, proliferation and differentiation into enteric neurons^2,3^. When RET is deficient, these progenitors fail to fully colonize the bowel, causing the congenital aganglionosis and functional bowel obstruction that are the cardinal features of this disorder^4^. The standard of care for HSCR is surgical resection of the aganglionic bowel segment. While surgery resolves the bowel obstruction, many HSCR patients continue to suffer from chronic defecatory dysfunction^5^. Deficient RET signaling in the non-resected, ganglionated bowel could play a role in these chronic complications; however, this possibility has not been well explored.

While RET has been studied extensively in fetal development, its functions in the postnatal intestine have received less attention. Identifying these functions is crucial, not just for understanding the chronic complications of HSCR, but also for addressing the prominent gastrointestinal (GI) symptomatology in the many settings in which RET signaling is disrupted. These include multiple endocrine neoplasia type 2 (MEN2), irritable bowel syndrome (IBS), and cancer chemotherapy with RET kinase inhibitors^6,7^. In the postnatal mouse intestine, there are at least 3 cell types with important roles in homeostasis that are potentially vulnerable to disruptions in RET signaling: immune cells, epithelial cells and neurons. In the immune system, RET signaling in group 3 innate lymphoid cells stimulates interleukin-22 secretion, which limits epithelial reactivity and vulnerability to inflammation^8^. In the epithelium, RET is expressed by isolated cells scattered throughout the intestine that colocalize with the enteroendocrine cell (EEC) marker chromogranin A (CHGA)^9,10^. EECs are diverse cells that release a variety of bioactive signals in response to chemical and mechanical cues from the gut lumen. Many EEC-derived signals alter intestinal motility including serotonin (5-HT). *In vitro*, RET ligands alter 5-HT secretion from colon organoids in a manner that is sensitive to RET kinase inhibition^9^. The significance of this signaling *in vivo*, and whether it plays a role in causing dysmotility in RET-deficient states like HSCR, however, are unclear.

Among the strongest candidates to mediate the effects of aberrant RET signaling on GI motility are enteric neurons, which are essential for normal gut motor function. While RET is necessary for the initial generation of neurons in the fetal ENS, it is rapidly downregulated in many of these neurons in both mice and humans^2,11^. The mature mammalian ENS is organized into two main ganglionated plexuses, myenteric and submucosal. In humans, RET can be detected in a subset of neurons in both plexuses of the adult duodenum and colon^12,13^. At least some of the *RET*^*+*^ colonic neurons contain transcripts for choline acetyltransferase (*CHAT*), indicating that they are cholinergic^14^. RET is also expressed in a subset of neurons in the adult mouse ENS^15^, but the identity of these neurons and whether RET serves any essential functions in the neurophysiology of the adult gut are undefined. Analogous to observations in EECs, the application of RET ligands to mouse enteric neurons *in vitro* alters neuropeptide secretion^16,17^. Similarly, RET ligand administration to gut explants changes calcium transients in some neurons and alters tissue contractility^18^; however, whether these effects are direct and how they relate to motor functions *in vivo* remain unknown. To better define the role of RET signaling in the postnatal intestine here we identify the major populations of RET-expressing neurons and EECs. Then, using conditional genetic approaches and gut-restricted RET kinase inhibition to disrupt RET signaling in one or more of these cell populations *in vivo*, we find that RET kinase activity in a subset of EECs limits peptide secretion to regulate small bowel motility.

## Materials and Methods

### Animals

See Supplementary Methods for mouse strain details. Littermate controls and 12-16 week old male and female mice were used for all experiments, except where noted. All studies were conducted in accordance with the GSK Policy on the Care, Welfare and Treatment of Laboratory Animals and were reviewed by the Institutional Animal Care and Use Committees at GSK, Columbia University Medical Center or Boston Children’s Hospital.

### Immunohistochemistry and quantitation

Intestinal tissue was prepared for immunostaining as previously described^19^. Primary and secondary antibodies used are listed in Supplemental Table 1. Methods for quantitation of CFP^+^ neurons and intestinal epithelial cells are detailed in Supplementary Methods. For quantitation of EECs in RET^EpiKO^ mice and controls, 12µm cryosections underwent IHC and were imaged by an investigator blinded to genotype. Only images with intact crypt-villus units, as determined by staining of cell nuclei, were used. At least 10 crypt-villus units were quantified per section per animal.

### Gastrointestinal motility testing and stool composition measurements

Measurements of gastrointestinal transit time (GITT), gastric emptying/small intestinal transit (GE/SITT), colonic migrating motor contractions (CMMCs), fecal pellet output (FPO), and fecal pellet composition were carried out as previously described^19^. To test effects of RET kinase inhibition, animals were gavaged twice daily at 8 AM and 6 PM with 10mg/kg GSK408B (an analog of a previously published RET inhibitor with poor systemic absorption^20,21^) or vehicle (0.5% hydroxypropylmethylcellulose) for 3.5-5.5 days. The final dose was administered 30min prior to GI motility testing. For standard GITT measurements, animals were gavaged with 300µL of 6% carmine red in 0.5% methylcellulose and placed in individual cages with *ad libitum* food and water. After 60min, cages were checked every 10 minutes for fecal pellets containing red dye. For fasting GITT measurements, food was removed 6 hours prior to dye gavage and water was available *ad libitum*. For 3 week-old mice, fecal pellets were checked starting 30 minutes after gavage of 100µL of 6% carmine red. To assess the effect of PYY, a single dose of BIIE0246 (Tocris 1700) or sterile water (vehicle) was administered via IP injection 30 minutes before GITT measurements. GITT, FPO, and GE/SITT measurements were initiated between 8:30-9am. For CMMC measurements, two colons were studied in tandem in the organ bath: one from a vehicle-treated animal and one from a drug-treated animal. All parameters were obtained as averages from three 15-minute videos analyzed by Scribble and Matlab scripts^22^. CMMCs were defined as beginning in the proximal colon and propagating > 50% of the colon length.

### Statistical analyses

Analyses were performed using SPSS and data were visualized using GraphPad Prism version 9. Two-tailed, unpaired t-tests were used to compare pairs of group means. For 3 or more experimental conditions, ANOVA was used to compare group means as detailed for each experiment in Supplemental Methods. Graphs show individual data points for each biological replicate (i.e. animal) along with mean and standard error of the mean (SEM), except where noted.

Please see Supplemental Methods for details of gene expression analysis by qPCR and scRNA-SEQ, quantitation of peptide hormones by immunoassays, and biogenic amines by liquid chromatography/mass spectrometry (LC/MS).

## Results

### Specific subsets of neurons in the adult ENS express RET

RET expression is maintained in at least some enteric neurons after fetal development^15^; however, the identity of the neurons and the extent to which RET expression varies along the radial and longitudinal axes of the ENS is unknown. To address these gaps, we characterized *Ret* expression in the postnatal ENS using whole-mount immunohistochemical staining of tissue from *Ret*^CFP/+^ knockin mice. The *Ret*^CFP/+^ genetic reporter enabled us to circumvent the limited efficacy of commercially available RET antibodies. In adult mice, *Ret*-expressing cells immunoreactive for the pan-neuronal marker ANNA-1 were abundant within the myenteric and submucosal plexuses (Figure 1A-C). In contrast, *Ret* was undetectable in intraganglionic and intramuscular glia marked by S100B immunoreactivity (Supplementary Figure 1). We quantified the proportions of *Ret*-expressing enteric neurons across 3 stages of the lifespan: pre-pubertal (4 weeks), mature adult (12-16 weeks) and aged adult (52 weeks). In the mature adult colon, 54.7% ± 6.1% of myenteric neurons and 69.2% ± 6.8% of submucosal neurons expressed *Ret* (mean ± SEM; n=8-9 mice/group). Age did not affect these proportions in either plexus (submucosal p=0.1274; myenteric p=0.1517); moreover, there were no differences in the proportions of *Ret*-expressing myenteric neurons between the colon and ileum (p=0.3317). These data indicate that *Ret* is expressed in a large subset of neurons in both plexuses of the postnatal mouse ENS and that the proportion remains relatively stable with age, with little variation along the length of the gut.

**Figure 1.**
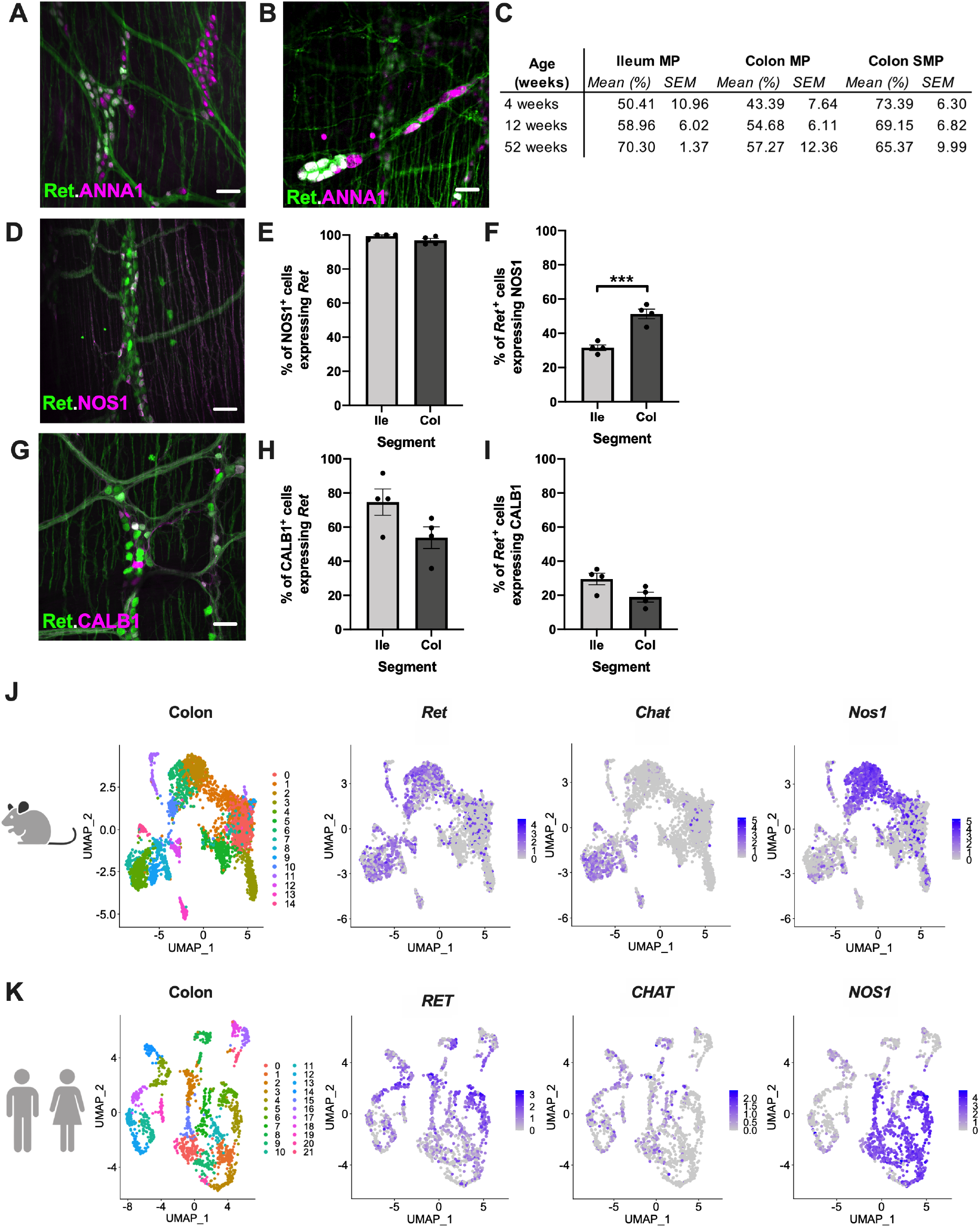
*Ret* is expressed in subsets of mature enteric neurons, including all NOS1^+^ nitrergic neurons and some CALB1^+^ cholinergic neurons. **(A-B)** *Ret* expression in the myenteric (A) and submucosal plexus (B) visualized by immunostaining adult *Ret*^CFP/+^ mouse colon for CFP and the pan-neuronal marker ANNA-1. **(C)** The proportions of ANNA-1^+^ neurons that expressed *Ret* (CFP^+^) in ileums and colons from *Ret*^CFP/+^ mice that were 4, 12-16, or 52 weeks of age in the myenteric (MP) and submucosal plexus (SMP) layers (n=5-9 mice). **(D)** Virtually all NOS1^+^ myenteric neurons in adult *Ret*^CFP/+^ mouse ileum expressed *Ret*. **(E)** Proportions of NOS1^+^ myenteric neurons expressing *Ret* in adult ileum (Ile) and colon (Col). **(F)** Proportions of *Ret*^*+*^ adult myenteric neurons that were NOS1^+^. *** p<0.001 by unpaired t-test **(G)** A subset of CALB1^+^ myenteric neurons in an adult *Ret*^CFP/+^ mouse colon expressed *Ret*. **(H)** Proportions of CALB1^+^ myenteric neurons expressing *Ret* in the adult intestine. **(I)** Proportions of *Ret*^*+*^ adult myenteric neurons that were CALB1^+^. **(J)** UMAPs of scRNA-SEQ of myenteric neurons from adult mouse colons^24^ show *Ret* expression within a large subset of *Chat*^*+*^ neurons and the majority of *Nos1*^*+*^ neurons. **(K)** UMAPs of scRNA-SEQ of myenteric neurons from adult human colons^25^ show similar distributions of *RET, CHAT* and *NOS1* expression across clusters. Scale bars=50µm

To determine whether *Ret* expression in enteric neurons was stochastic or a fixed feature of specific subtypes, we further characterized the *Ret*^*+*^ myenteric neurons in mature adult *Ret*^CFP/+^ mice. In the myenteric plexus, nitrergic and cholinergic neurons are considered mutually exclusive populations with developmentally distinct trajectories^23^. Almost 100% of NOS1-immunoreactive neurons expressed *Ret* in the ileum and colon, suggesting that *Ret* expression is a fixed feature of nitrergic neurons (Figure 1D-E). NOS1^+^ neurons constituted 31.6% ± 1.6% of all *Ret*^*+*^ neurons in the ileum and 51.4% ± 2.8% in the colon (Figure 1F). In contrast, only a subset of myenteric neurons immunoreactive for Calbindin (CALB1), a calcium binding protein that marks many cholinergic neurons, expressed *Ret* (Figure 1G). In the ileum and colon, 74.7% ± 7.8% and 53.9% ± 6.4% of CALB1^+^ neurons expressed *Ret*, respectively (Figure 1H). CALB1^+^ neurons constituted a minority of *Ret*^*+*^ neurons, 29.6% ± 3.4% in the ileum and 19.0% ± 2.9% in the colon (Figure 1I). Analysis of *Ret* expression in published single cell transcriptional profiling data of myenteric neurons in adult mice was consistent with our observations that *Ret* expression is enriched in nitrergic neurons and subsets of cholinergic and other neurons^24^ (Figure 1J; Supplemental Figure 2). This expression pattern was similar in human colonic myenteric neurons^25^ (Figure 1K; Supplemental Figure 2), suggesting that RET signaling is likely to have conserved functions in the adult mammalian ENS.

### Genetic depletion of RET in adult mice transiently slows GI motility

RET is required for ENS development and *Ret*-null mice do not survive beyond birth because of this and other developmental defects^3^. Even mice lacking RET only in neural crest derivatives do not survive to adulthood due to intestinal aganglionosis^26^. Thus, the functions of RET in the mature ENS have remained unclear. To determine these functions, we generated *Ret*^CreER/flox^ mice (hereafter RET^cKO^) in which RET could be conditionally depleted in adult mice after ENS development. At baseline, RET^cKO^ mice are haploinsufficient for *Ret*. Upon administration of tamoxifen to induce Cre recombinase activity, cells that express *Ret* at the time of induction become *Ret*-null (Figure 2A). To determine the effects of postnatal RET depletion, adult RET^cKO^ mice and controls were induced with tamoxifen and studied for 4 weeks. RET^cKO^ mice appeared well and had no weight loss (Supplementary Figure 3A). Previous work showed that conditional deletion of RET in embryonic development after colonization of the bowel by ENS precursors caused colonic hypoganglionosis associated with cell death^27^, suggesting that enteric neurons require RET for survival. To test this possibility, we quantified neurons in the small and large intestines of RET^cKO^ mice and found that overall neuronal density (Figure 2B) and the proportions of NOS1^+^ and CALB1^+^ neurons (Supplemental Figure 3B) did not differ from those of littermates with intact *Ret* (*Ret*^flox/+^ or *Ret*^flox/flox^ mice), indicating that RET signaling is not essential for the survival of mature enteric neurons. The colonic hypoganglionosis previously reported with *Ret* ablation after ENS colonization may be attributable to a requirement for *Ret* in precursors that enter the bowel later via extrinsic nerves^28^.

**Figure 2.**
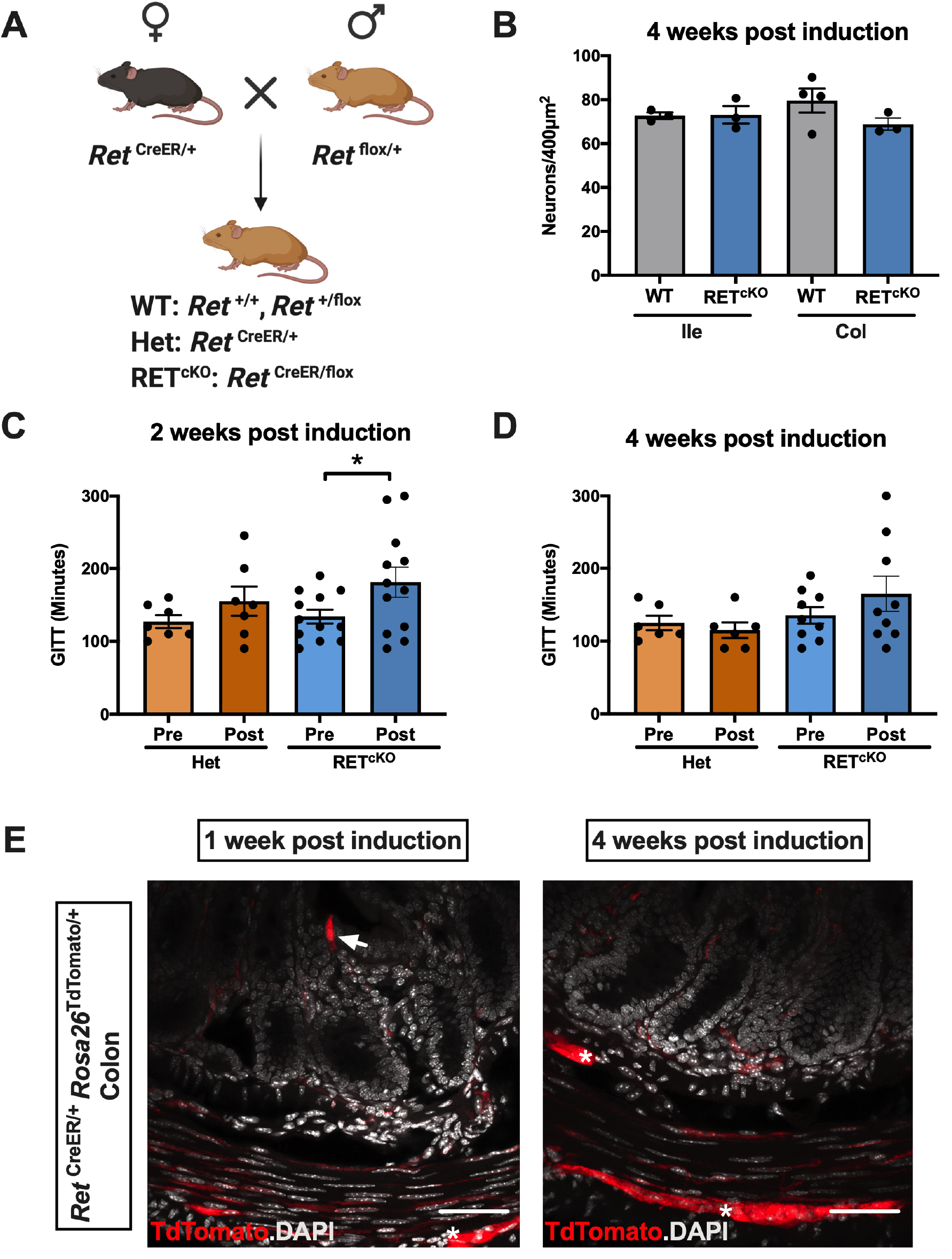
Conditional deletion of RET in adult mice causes transient GI dysmotility. **(A)** Strategy for generation of adult, conditional RET knockout mice (*Ret*^CreER/flox^; referred to as RET^cKO^) and two types of littermate controls (*Ret*^flox/+^ and *Ret*^flox/flox^; WT) and *Ret*-haploinsufficient mice (*Ret*^CreER/+^; Het). **(B)** Postnatal RET depletion does not alter myenteric neuronal density in the ileum or colon. **(C-D)** RET depletion slows GI transit time (GITT) in RET^cKO^ mice, but not in Het controls, at 2 weeks post-induction. GITT begins to normalize by 4 weeks. * p<0.05 by two-way repeated measures ANOVA **(E)** Colon cross-sections from *Ret*^CreER/+^ *Rosa26*^tdTomato/+^ mice 1- or 4-weeks after induction with tamoxifen. DAPI marks cell nuclei (white) and TdTomato (red) marks cells that either expressed *Ret* at the time of induction or derived from cells expressing *Ret* at that time. At 1-week, TdTomato is evident in enteric neuronal fibers and ganglia (*), as well as isolated epithelial cells (white arrow). At 4 weeks, neuronal TdTomato remains evident but epithelial expression is undetectable.

In the adult ENS, *Ret* was expressed in at least two populations of myenteric neurons that play important roles in peristalsis: NOS1^+^ inhibitory motor neurons, and CALB1^+^ neurons, many of which are intrinsic primary afferents^29,30^. To determine whether RET depletion affected the function rather than the survival of these neurons, we measured gastrointestinal transit times (GITT) in RET^cKO^ mice at 2- and 4-weeks post induction. To account for potential effects of *Ret* haploinsufficiency, we compared them to *Ret*^CreER/+^ littermates. Although GITT in control mice was no different at 2 or 4-weeks post-induction compared to baseline, GITT in RET^cKO^ mice was significantly prolonged at 2 weeks (Figure 2C). This defect resolved by 4-weeks (Figure 2D), suggesting that RET depletion in adult mice causes transient dysmotility.

Some enteric neurons may be continuously replaced in the adult ENS^31^. To determine whether turnover of RET-expressing neurons explained the transient dysmotility in RET^cKO^ mice, we generated *Ret*^CreER/+^*Rosa26*^TdTomato/+^ mice in which *Ret*-expressing cells could be permanently labeled with the TdTomato reporter. We administered tamoxifen to adult *Ret*^CreER/+^*Rosa26*^TdTomato/+^ mice to induce reporter expression and examined the small and large intestines at 1- and 4-weeks post-induction. At 1-week, robust TdTomato was evident in both plexuses of the ENS, immune cells in Peyer’s patches, and isolated epithelial cells (Figure 2E; Supplementary Figure 4). At 4-weeks, TdTomato expression remained unchanged in the neuronal and immune compartments but was no longer detectable in the epithelium (Figure 2E; Supplementary Figure 4). These observations suggest that RET-expressing epithelial cells turned over within 4-weeks while most RET-expressing neurons did not. RET^cKO^ mice, by proxy, are likely to have evolving levels of RET signaling in the gut as RET-expressing cell types turn over at different rates. Normalization of dysmotility in RET^cKO^ mice by 4-weeks post-induction could thus reflect repopulation of the intestinal epithelium with cells containing intact *Ret* alleles, incomplete RET depletion in the ENS, or mobilization of compensatory mechanisms within RET-expressing neurons. To distinguish between these possibilities and establish a stable model of RET deficiency in the mature ENS, we examined the effects of constitutively depleting RET in a population of differentiated enteric neurons.

Given the uniform expression of RET among inhibitory motor neurons and their well-established role in GI motility, we generated mice constitutively lacking RET in this population. Inhibitory motor neurons are characterized by expression of both VIP and NOS1, and most NOS1-expressing cells in the murine myenteric plexus express VIP^18,23,29,30^. Because *Nos1*^Cre/+^ mice, but not *Vip*^Cre/+^ mice, exhibited extensive Cre activity in the gut epithelium in addition to the ENS, we generated *Vip*^Cre/+^*Ret*^flox/flox^ mice to determine the effects of depleting RET expression within inhibitory motor neurons (hereafter RET^VipKO^). RET^VipKO^ mice were born at expected Mendelian ratios, appeared healthy, and grew at a pace similar to that of Cre-negative *Ret*^flox/flox^ littermates (Supplementary Figure 5A-B). As we observed with adult RET^cKO^ mice, constitutive depletion of RET in inhibitory motor neurons did not grossly alter the appearance or proportions of these neurons (Supplementary Figure 5C-D), again indicating that RET signaling was not essential for their survival. Surprisingly, adult RET^VipKO^ mice showed no deficits in GITT or 1-hour FPO (Supplementary Figure 5E-F), implying that RET signaling in this large population of motor neurons was not essential for gut motility. Unlike NOS1, VIP is also expressed in a large subset of submucosal neurons that can play a role in fluid flux across the epithelium. RET^VipKO^ mice exhibited no deficits in stool water content or fecal pellet size (Supplementary Figure 5G-H), leaving the functional significance of RET expression in VIP neurons to be determined. Taken together, our observations indicate that RET depletion in adult mice causes transient GI dysmotility. This dysmotility was not replicated by constitutively deleting RET in a large population of enteric neurons that are important for motility, suggesting that RET expression in epithelial cells may be more consequential for propulsive activity.

### RET signaling in the gut epithelium regulates motility selectively in male mice

To determine whether epithelial RET signaling was necessary for normal GI motility, we generated *Vil1*^Cre/1000^ *Ret*^flox/flox^ mice (hereafter RET^EpiKO^) that would lack RET in the intestinal epithelium but not in the ENS^32^. RET^EpiKO^ mice were born at expected Mendelian ratios, had no overt deficits, and grew at the same pace as their littermate controls (*Ret*^flox/flox^ mice; Supplementary Figure 6A-B). *Ret* transcripts were undetectable in the epithelium of RET^EpiKO^ mice although expression in the non-epithelial compartment remained robust (Supplementary Figure 6C). In contrast to RET^VipKO^ mice, GI transit was 30% slower in adult RET^EpiKO^ mice than controls; however, this deficit was only present in males (Figure 3A). Both male and female RET^EpiKO^ mice exhibited no alterations in FPO (Figure 3B), stool water content or fecal pellet mass (Supplementary Fig 6D and E), all of which are primarily measures of colonic function. These observations establish that epithelial RET signaling is essential for normal GI motility in males and suggest that its effects are most prominent in the upper GI tract.

**Figure 3.**
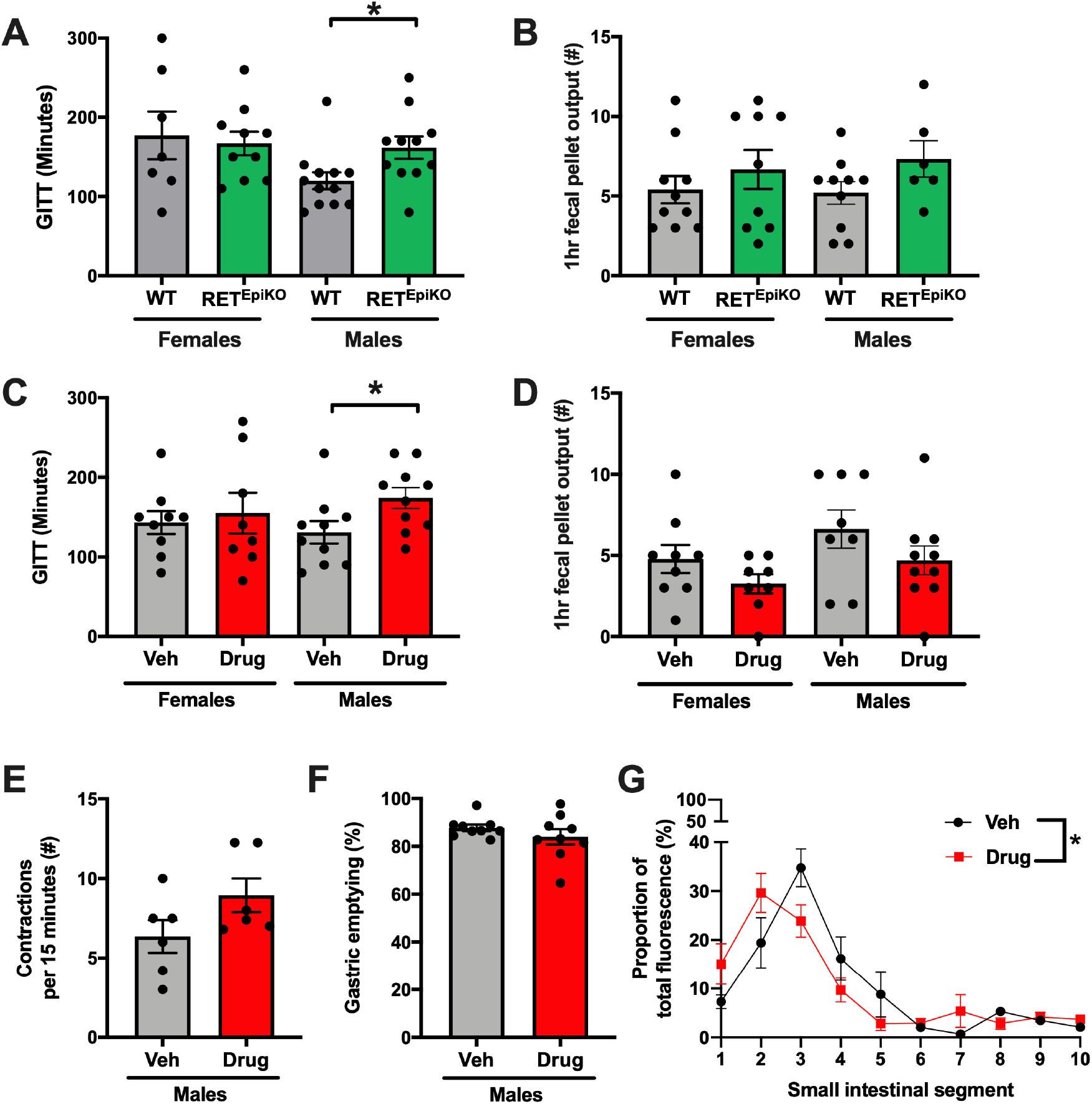
Genetic or pharmacologic disruption of intestinal epithelial RET signaling slows GI transit in a sex-dependent manner. **(A)** GI transit time in adult mice lacking RET in the intestinal epithelium (*Vil1*^Cre^ *Ret*^flox/flox^; RET^EpiKO^) is 30% slower in males, but not females, compared to controls (*Ret*^flox/flox^; WT). **(B)** 1-hour FPO was no different in RET^EpiKO^ mice compared to controls. **(C)** GI transit time in mice administered a gut-restricted RET kinase inhibitor (Drug) was 30% slower in males, but not females, compared to vehicle-treated controls (Veh). **(D)** RET inhibition did not alter 1-hour fecal output in either sex. **(E-F)** Colonic contraction frequency and gastric emptying were unaffected by RET inhibition. **(G)** Small intestinal transit, measured as the distance traveled by a fluorescent dye 15 minutes after orogastric gavage, was slowed by RET inhibition. The x-axis represents intestinal segment number (proximal?distal; n=8-9 mice). * indicates p<0.05 by two-way ANOVA (**A-D**) or one-way repeated measures ANOVA (**G**).

To ascertain whether acute disruption of epithelial RET signaling would have the same effects as chronic genetic depletion, we determined the effects of RET kinase inhibition on GI motility. GSK408B belongs to a family of RET kinase inhibitors that are highly specific to RET kinase and have poor systemic absorption, such that enteral administration enables drug activity to be gut-restricted^21^. We administered GSK408B or vehicle to wildtype mice twice daily for 3.5 days by orogastric gavage and then assessed GI motility. Remarkably, RET kinase inhibition phenocopied RET^EpiKO^ mice. Male, but not female, mice displayed 30% longer GI transit times with no difference in 1-hour FPO (Figure 3C-D). To delineate which segments of the GI tract relied on RET signaling for normal motility, we examined gastric emptying, small intestinal transit and colonic motility in GSK408B-treated males. Consistent with normal colonic function seen by 1-hour FPO measurements *in vivo*, detailed analysis of colonic migrating motor contractions *ex vivo* did not reveal a difference in contraction frequency or any other feature of these motor behaviors (Figure 3E; Supplemental Figure 6F-G). In the upper GI tract, RET kinase inhibition did not alter gastric emptying but did slow small intestinal transit (Figure 3F-G). In conclusion, both pharmacological and genetic disruption of RET kinase slowed GI motility selectively in males with effects primarily manifested on the small intestine.

The sex-dependent effects of disrupting epithelial RET were striking and resembled the sex-dependent penetrance of RET mutations, a major factor in the disproportionate incidence of HSCR in males. The role of epithelial RET in motility may similarly be established during fetal development or acquired later, such as upon puberty, when gonadal steroids surge and influence gut motility^22^. To distinguish between these possibilities, we assessed GI motility in 3-week-old, pre-pubertal RET^EpiKO^ mice. At this age, epithelial RET depletion had no effect on GI motility in males (Supplementary Figure 7), supporting the idea that RET-dependence of motility in males is an acquired rather than developmental effect.

### Subsets of enteroendocrine cells in the adult intestinal epithelium express *Ret*

To determine how epithelial RET signaling promotes motility, we examined RET expression in the adult small intestinal epithelium. EECs are sensory epithelial cells that detect chemical and mechanical stimuli from the gut lumen; at least some express RET^9,10^. Consistent with these reports, in 12-16 week old *Ret*^CFP/+^ mice, the majority of *Ret*-expressing cells in the duodenum and ileum were immunoreactive for the EEC marker CHGA in both sexes (Figure 4A-C). Similar to enteric neurons, *Ret* expression was not universal among EECs. In the male duodenum, for example, 49% ± 6% of ChgA^+^ cells expressed *Ret*, suggesting that *Ret* expression might be characteristic of specific types of EECs.

**Figure 4.**
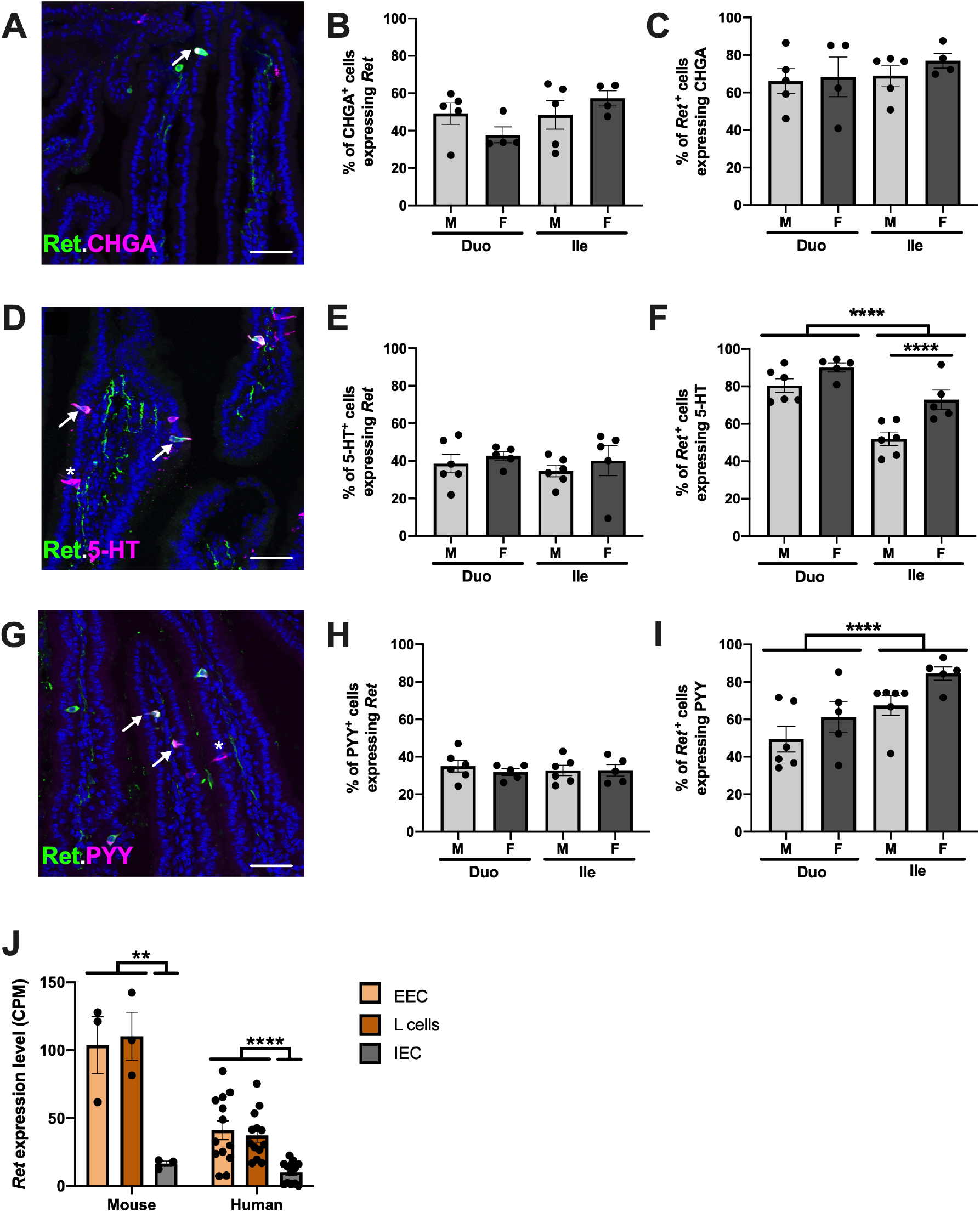
The majority of *Ret*-expressing small intestinal epithelial cells are enteroendocrine cells, including subsets of enterochromaffin and L-cells. **(A)** *Ret*-expressing cells are immunoreactive for CHGA in the *Ret*^CFP/+^ mouse small intestine. **(B)** Large subsets of CHGA^+^ epithelial cells in adult male (M) and female (F) *Ret*^CFP/+^ mice express *Ret*, in the duodenum (Duo) and ileum (Ile). **(C)** The majority of *Ret*-expressing cells in the epithelium express CHGA. **(D)** Many small intestinal epithelial cells immunoreactive for serotonin (5-HT) express *Ret* (arrows) in a *Ret*^CFP/+^ mouse; but some 5-HT^+^ cells are *Ret*-negative (*). **(E)** Subsets of enterochromaffin cells throughout the small intestine express *Ret*. **(F)** Most *Ret*-expressing cells in the intestinal epithelium contain 5-HT. The proportion was higher in the duodenum than the ileum in both sexes. **(G)** Many small intestinal epithelial cells immunoreactive for PYY express *Ret* (arrows) in a *Ret*^CFP/+^ mouse; but some PYY^+^ L-cells are *Ret*-negative (*). **(H)** A subset of PYY^+^ cells expresses *Ret* throughout the small intestine in both sexes. **(I)** Many *Ret*-expressing small intestinal epithelial cells contain PYY. The proportion of *Ret*^*+*^ cells that are L-cells is higher in the ileum and is no different between the sexes. **(J)** Analysis of *Ret* expression in transcriptional profiling data^33^ shows that *Ret* is highly expressed in enteroendocrine cells (EEC) overall, as well as L-cells in particular, compared to other intestinal epithelial cells (IEC), in both mice and humans. * p<0.05, ** p<0.01, and **** p<0.0001 by two-way ANOVA (**A-I**) or one-way ANOVA (**J**).

EECs are diverse cells that are often classified by the biogenic amines and peptides they release in response to stimuli. Two major types of EECs that regulate GI motility are enterochromaffin cells and L-cells, which release 5-HT and PYY, respectively. In the male duodenum, 38.5% ± 4.9% of enterochromaffin cells marked by 5-HT immunoreactivity expressed *Ret*, and this proportion was consistent across both sexes and in the ileum (Figure 4D-E). Interestingly, while 80-90% of *Ret*^+^ cells expressed 5-HT in the duodenum, this proportion was lower in both sexes in the ileum (Figure 4F), highlighting the potential for regional differences in epithelial RET biology. In terms of L-cells, 35% ± 3.2% of PYY^+^ epithelial cells in the male duodenum expressed *Ret*, and this proportion was consistent across both sexes and along the length of the small intestine (Figure 4G-H). Conversely, 49.4% ± 6.9% of *Ret*^+^ cells expressed PYY in the male duodenum, and this proportion was even higher in the ileum (Figure 4I). Secondary analysis of transcriptional profiling data^33^ from small intestinal EECs, L-cells and intestinal epithelial cells (IEC) further showed that *Ret* expression was highly enriched in EECs and L-cells in both mice and humans (Figure 4J). In sum, *Ret* is expressed in large subsets of enterochromaffin cells and L-cells, two major types of EECs that influence GI motility.

### RET signaling alters GI motility by dampening peptide hormone release

Enterochromaffin cells respond to mucosal deformation by releasing 5-HT onto local nerve endings to stimulate peristalsis^34^. While the necessity of enterochromaffin-derived 5-HT for motility has been debated, genetic elimination of these cells or disruption of their mechanosensory apparatus clearly slows GI transit^35,36^. Enterochromaffin cells are also a major source of circulating 5-HT and loss of 60-70% of these cells is sufficient to dramatically reduce both serum and intestinal levels^36^. To determine whether disrupting RET signaling in the gut slowed motility in adult males by compromising enterochromaffin cell-derived 5-HT, we administered the RET kinase inhibitor GSK408B to male mice for 3.5 days and measured fasting and post-prandial levels of 5-HT and its breakdown product 5-HIAA. RET kinase inhibition had no effect on 5-HT or 5-HIAA levels in the blood or intestinal tissues (Supplementary Figure 8), suggesting that disrupting RET signaling in the adult gut does not slow motility by compromising 5-HT secretion.

Unlike enterochromaffin cells which respond to both mechanical and chemical stimuli, L-cells are generally considered chemosensory cells that release peptides in response to luminal nutrients^37^. These peptides, such as PYY and glucagon-like peptide 1 (GLP1), signal by endocrine, paracrine and neurocrine mechanisms to influence a variety of gut functions including motility. Although RET kinase inhibition had no effect on serum 5-HT, it was associated with much higher levels of GLP1 and PYY in the post-prandial state (Figure 5A-B). These elevated levels were evident in males, but not females, similar to the finding of slowed GI transit, suggesting that the phenotypes might be linked.

**Figure 5.**
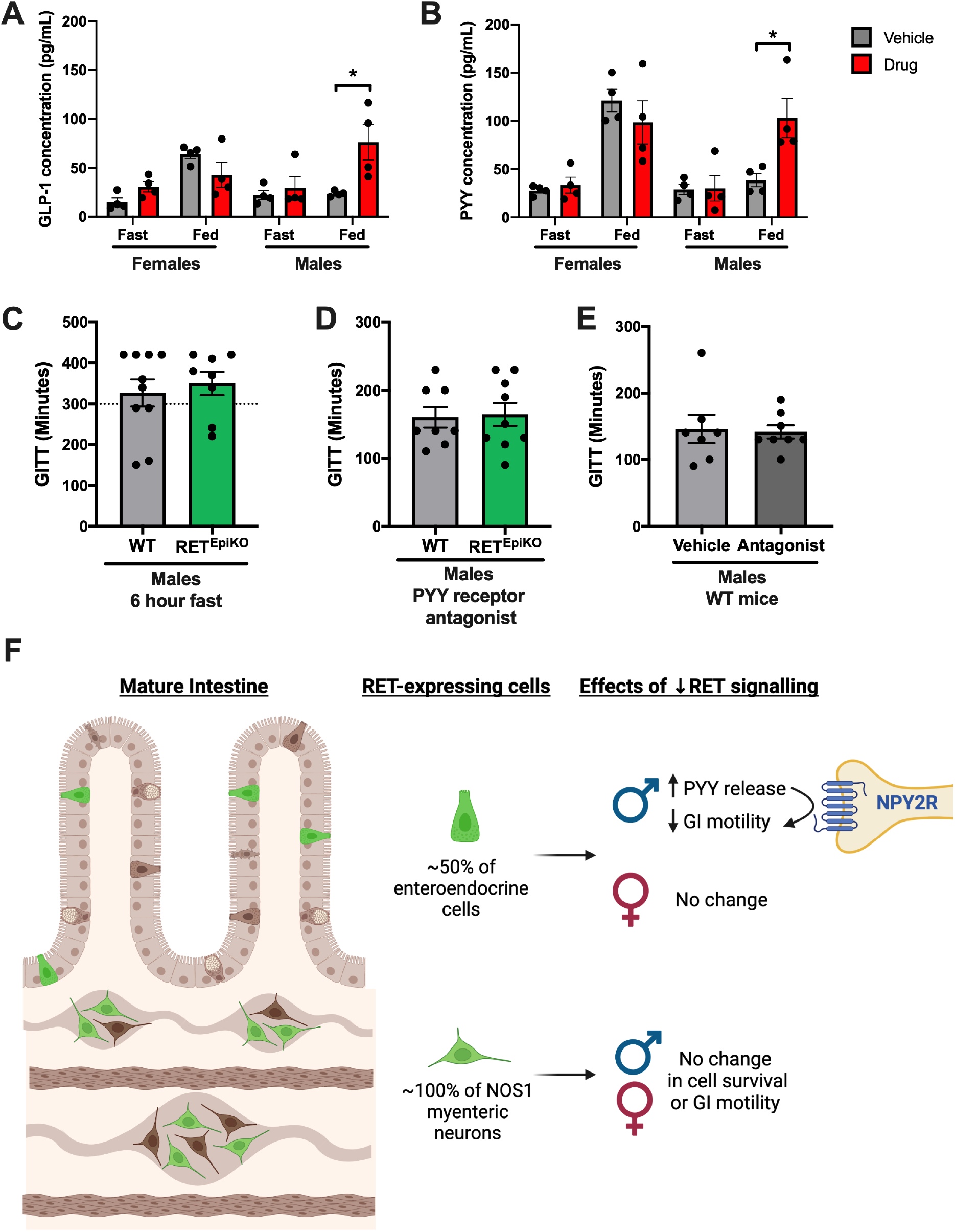
Disruption of RET signaling leads to exaggerated PYY release, slowing GI motility in a sex- and nutrient-dependent manner. **(A-B)** Serum levels of GLP1 and PYY after overnight fast (Fast) or 2 hours post re-feeding (Fed) in mice administered a RET kinase inhibitor (Drug) or vehicle (Veh). In females, GLP1 and PYY rise with feeding, as expected, and are no different between RET inhibitor and vehicle-treated mice. In males, fasting levels of GLP1 and PYY levels are low in both conditions, but feeding leads to much higher levels of both peptide hormones in RET inhibitor-treated mice. * indicates p<0.05 by two-way ANOVA. **(C)** Male RET^EPIKO^ mice have the same average GI transit times as controls (WT) following a 6-hour fast. Dotted line indicates the cut-off time for the standard, non-fasting GITT assay. **(D)** A single dose of the NPY2R antagonist BIIE0246 restores GITT in male RET^EPIKO^ mice to that of controls (see Figure 3A for comparison). **(E)** A single dose of BIIE0246 does not alter GITT in WT mice. **(F)** A model of RET expression and proposed functions in the mature intestine.

L-cells normally secrete GLP1 and PYY in a nutrient-dependent manner. To determine whether the dysmotility associated with epithelial RET deficiency was nutrient-dependent, we measured GI transit in male RET^EpiKO^ and littermate controls after a 6-hour fast. Fasting slowed GITT among all mice, as expected, but also eliminated the effect of RET deficiency; indicating that dysmotility in RET^EpiKO^ mice was nutrient-dependent (Figure 5C). PYY mediates the ileal brake phenomenon, in which nutrient-stimulated release of PYY from L-cells slows upper GI motility^38^. PYY signals through two different receptors, NPY1R and NPY2R. Upon secretion from L-cells, PYY is cleaved into an active form (PYY3-36) that has highest affinity for NPY2R; NPY2R signaling has been shown to modulate GI motility^39^. To determine whether exaggerated PYY signaling was responsible for slowed GI transit in male RET^EpiKO^ mice, we administered a single dose of the NPY2R antagonist BIIE0246 to RET^EpiKO^ males and littermate controls prior to GITT measurement. NPY2R inhibition was sufficient to completely rescue the motility deficit in RET^EpiKO^ mice (Figure 5D) at a dose that had no effect on motility in wildtype males (Figure 5E), suggesting that excessive PYY signaling was necessary for slowing GI motility in RET-deficient males. This excessive signaling could occur if supernumerary L-cells are present in the absence of RET or if RET signaling normally limits the production or release of PYY. RET^EpiKO^ mice had similar L-cell numbers and *Pyy* transcript levels as controls (Supplementary Figure 9), suggesting that RET most likely affects PYY release. Taken together, these observations support the conclusion that RET signaling in L-cells stimulates intestinal motility in males by limiting the amount of PYY that is released and available to act on NPY2R^+^ neurons following a meal (Figure 5F).

## Discussion

RET is a potent receptor tyrosine kinase that plays myriad roles in development but its homeostatic functions in the mature intestine have remained poorly explored. Here, we report that important subsets of enteric neurons and EECs in the adult mouse and human intestines express RET. Using genetic and pharmacologic approaches to disrupt RET signaling in the adult intestine, we discovered that RET activity in EECs, but not neurons, stimulates intestinal motility by regulating post-prandial PYY release. Unexpectedly, this RET-dependent effect on motility was sex-specific.

*Ret* expression in the fetal mouse and human ENS becomes restricted to one of two major developmental trajectories for enteric neurons^11,23^. Our study extends these observations to the adult ENS, demonstrating that *Ret* expression is stably maintained by 55-70% of neurons in the myenteric and submucosal plexuses throughout life. Although mature cholinergic neurons vary in *Ret* expression, NOS1^+^ neurons in the myenteric plexus, most of which are VIP^+^ inhibitory motor neurons, virtually all express *Ret*. We constitutively depleted RET in these neurons, however, and did not uncover any deficits in GI motility or cell survival. More detailed analyses of gut motor behaviors may be necessary to reveal deficits; alternatively, RET signaling in adult enteric neurons may be more important in the context of injury than homeostasis. Given the long-term risk of enterocolitis in patients with HSCR, this possibility is important to explore. Transcriptional profiling studies have shown robust RET co-receptor expression among enteric glia^18,25^; however, we found no evidence of RET expression in intramuscular or myenteric glia. Gut lymphoid cells that express RET co-receptors, but not RET, signal *in trans* with neighboring RET^+^ cells^40^. If enteric glia participate in RET signaling beyond ligand secretion, they may similarly signal *in trans* with neighboring RET^+^ neurons.

In contrast to our findings in the ENS, three independent approaches to RET depletion in the intestinal epithelium revealed an essential role for RET in EEC regulation of GI motility. First, we found that conditional deletion of all RET signaling in adult *Ret*^CreER/flox^ mice led to transient dysmotility that resolved as the gut epithelium turned over. The time course of this defect was similar to the transient dysmotility that occurs upon enterochromaffin cell depletion, which resolves as the cells are restored^36^. Second, we found that constitutive RET depletion selectively in the intestinal epithelium slowed GI transit by 30% in male mice. Third, pharmacologic inhibition of RET kinase in adult mice phenocopied the genetic models of RET depletion. These observations converge to support the conclusion that RET signaling in EECs is necessary for regulating intestinal motility in males.

Although subsets of both enterochromaffin cells and L-cells expressed RET in the adult intestine, we found that RET kinase inhibition *in vivo* altered levels of PYY and GLP1, but not 5-HT. EECs secrete many signals and our observations suggest selectivity of RET effects. Further work will be needed to delineate this selectivity and determine how RET signaling modulates EEC secretion. Recent studies have shown that EECs engage in direct synapse-like communication with vagal afferents via cellular extensions called neuropods^41^. RET effects on this type of communication remain to be determined. EECs express multiple RET co-receptors in both mice and humans, and extend projections in response to a RET ligand *in vitro*^33,41^. It will be interesting to determine the identity and cell sources of the relevant ligands, and whether the effects of RET inhibition in EECs are ligand-specific.

Even though male and female EECs expressed RET, RET depletion or inhibition only perturbed GI motility and hormone release in males. This observation resembles the striking sexual dimorphism of human HSCR, which occurs 4 times more often in males than females^4^. GI motility was intact in juvenile RET^EpiKO^ males, suggesting that puberty alters male EECs and/or their cell circuits. Gonadal androgen signaling to the ENS regulates colonic motility in post-pubertal males^22^. Although there is no evidence for androgen effects on EECs, estrogen has been shown to stimulate endocrine cell secretion^42^. And *in vitro*, human female EECs exhibit lower thresholds for nutrient-dependent 5-HT release than male EECs^43^. These features of EEC biology may converge to make male EECs more vulnerable to the effects of RET depletion, leading to the sex-dependent phenotypes observed. More broadly, our observations add to the accumulating evidence that the cellular-molecular mechanisms governing gut motility are sex-dependent.

Overall, our findings have implications for the many clinical contexts in which RET signaling is disrupted, including HSCR, MEN2, and cancer chemotherapy targeting RET kinase. Perturbations of RET signaling in EECs could explain chronic GI dysmotility in HSCR as well as the high rates of diarrhea and constipation observed in clinical trials with RET kinase inhibitors^6^. Conversely, this nutrient-dependent mechanism could be exploited to treat post-prandial GI symptoms in patients with functional GI disorders. Gut-restricted RET kinase inhibitors are already in clinical trials for IBS and are highly effective in pre-clinical models of visceral hypersensitivity^20^. Some anti-nociceptive effects of these compounds may be attributable to their effects on EECs, which directly communicate with visceral afferent neurons^44^. In conclusion, we have found that *Ret* expression is prominent in the adult ENS and intestinal epithelium of both mice and humans. Disruption of epithelial RET kinase signaling in mice revealed an essential, sex-dependent function of RET in EECs that impacted their regulation of intestinal motility. These observations establish a role for RET signaling in GI homeostasis *in vivo* that could be targeted for the treatment of digestive disorders.

## Supporting information

Shepherd_et_al_2022_Supplementary_Materials

## Abbreviations

CFP: cyan fluorescent protein
CMMC: colonic migrating motor complex
dpt: days post treatment
EEC: enteroendocrine cell
ENS: enteric nervous system
FPO: fecal pellet output
GE/SITT: gastric emptying/small intestinal transit time
GI: gastrointestinal
GITT: gastrointestinal transit time
HSCR: Hirschsprung’s Disease
IBS: irritable bowel syndrome
IPAN: intrinsic primary afferent neuron
LC/MS: liquid chromatography/mass spectrometry
MEN2: multiple endocrine neoplasia type 2
MP: myenteric plexus
PYY: peptide YY
scRNA-SEQ: single cell RNA sequencing
SMP: submucosal plexus
UMAP: uniform manifold approximation and projection
WT: wildtype
5-HT: serotonin

## Author Contributions

A.S., L.F., S.S., D. R., E. M., M.D.G, and M.R. conceived aspects of the initial study. A.S., L.F., S.S., D.R., E.M., L. M., A.R. and M.R. performed experiments and analyzed results. A.M. performed single cell RNA sequencing analysis. M.R. and M.D.G. obtained funding for the studies. A.S. and M.R. wrote the manuscript. All authors edited and approved the final manuscript.

## Acknowledgments

We are grateful to V. Lennon (Mayo Clinic) for ANNA-1 antisera, H. Enomoto (Kobe University) for *Ret*^CFP/+^ mice, D. Ginty (Harvard Medical School) for *Ret*^flox/flox^ and *Ret*^CreER/+^ mice, and members of the Rao laboratory for helpful discussions and experimental support. We thank J. Russell and S. Kumar at GlaxoSmithKline for providing GSK408B and helpful discussions. We thank D. Breault and M. Rutlin for critical reading of the manuscript. Schematics in Figures 2A and 5F were created with BioRender.com. This study received partial funding support from sponsored research agreements with GlaxoSmithKline and Boston Pharmaceuticals, as well as funding from the Schmidt Science fellowship (A.S.), NIH T32DK091227 (in support of L.F.), NSF graduate fellowship (A.M.), NIH R01NS15547 (M.D.G), philanthropic support from Ivan and Phyllis Seidenberg (M.R.), Paul Marks Scholar Award (M.R.), Smith Family Foundation Odyssey Award (M.R.) and NIH K08DK125636 (M.R.). Core facilities utilized were supported by the Harvard Digestive Disease Center (NIH P30DK034854) and the Boston Children’s Hospital/Harvard Medical School Intellectual and Developmental Disabilities Research Center (NIH U54 HD090255).

