## Supplementary material for "RET Signaling Persists in the Adult Intestine and Stimulates Motility by Limiting PYY Release from Enteroendocrine Cells": Shepherd_et_al_2022_Supplementary_Materials

#### Supplementary Methods

#### Animals

*Rosa26*$^{TdTomato/TdTomato}$ (Strain 007909), FVB/NJ (Strain 001800), Vil1$^{Cre/1000}$ (Strain 021504), and *Vip*$^{Cre/+}$ mice (Strain 010908) were purchased from Jackson Laboratories. *Ret*$^{CFP/+}$ mice (MGI: 3777556) were generously donated by Hideki Enomoto (Kobe University). *Ret*$^{CreER/+}$ (MGI: 4437245) and *Ret*$^{flox/flox}$ (MGI: 5698695) mice were generously donated by David Ginty (Harvard Medical School). All transgenic mice were on a C57BL/6J background except *Ret*$^{flox/flox}$, which were on a mixed C57BL/6J-FVB/NJ background to optimize breeding. *Ret*$^{CreER/flox}$, RET$^{EpiKO}$ and RET$^{VipKO}$ mice were generated from respective parental strains. *Ret*$^{CreER/flox}$ mice, *Ret*$^{CreER/+}$*Rosa26*$^{TdTomato/+}$ mice, and controls were administered 8mg of tamoxifen by orogastric gavage at 12-16 weeks of age once daily for 1-3 doses and then studied at 7, 14 or 28-30 days post-treatment (dpt). All mice were housed in specific pathogen-free (SPF) facilities at Columbia University Medical Center or Boston Children's Hospital on a 12 hour light-dark cycle.

#### Quantitation of immunohistochemical staining

For neuronal quantitation, intestinal segments from *Ret*$^{CFP/+}$ mice were subjected to whole-mount immunostaining for CFP and molecular markers of interest, and then optically cleared. Single planar images were obtained on a Nikon A1R confocal microscope of 6 fields (myenteric plexus) or 10-12 fields (submucosal plexus) per segment per animal at 12.5X magnification. Images were randomized and analyzed either by an automated protocol on Volocity (PerkinElmer) for the myenteric plexus[19], or quantified manually on ImageJ for the submucosal plexus. Figure 1 shows representative images of gut segments from 12-16 week old *Ret*$^{CFP/+}$ mice immunostained as whole mounts and imaged at the level of the myenteric plexus.

For *Ret*-expressing epithelial cell quantitation, 12µm cryosections of intestine from *Ret*$^{CFP/+}$ mice were immunostained for CFP and the marker of interest. Twelve images per segment per animal were captured using an Olympus epifluorescence microscope at a magnification of 10X, and then manually quantified with ImageJ using the CellCounter plugin. The total number of CFP$^{+}$ or marker$^{+}$ epithelial cells was first scored independently and then the proportion with colocalization was calculated. Antibody information is available in Supplementary Table 1. Figure 4 shows representative images of frozen sections from small intestines isolated from 12-16 week old *Ret*$^{CFP/+}$ mice immunostained and imaged at the level of the villi.

### Gene expression analysis

For *Ret* quantitation in WT mice, intestinal segments from 3-week-old or 12-week-old male and female FVB/NJ mice were washed with chilled PBS and then subjected to mechanical scraping to isolate the epithelial layer. For *Ret* and *Pyy* quantification in male $Ret^{EpiKO}$ mice, the same procedure was used with the addition of a brief pre-incubation in 5mM EDTA prior to scraping. Epithelia were collected into Trizol (Invitrogen) for RNA extraction. RNA was purified through RNeasy columns (Qiagen) and converted to cDNA (iScript cDNA Synthesis kit, BioRad). Quantitative RT-PCR was done with SYBR Green Select Master Mix (Thermo Fisher). Primer sequences are listed in Supplementary Table 2. *Ret* expression in sorted EEC, L-cells and intestinal epithelial cells from mice and humans was assessed in a published dataset[33]. Raw data were downloaded from GREIN[45] using Gene Expression Omnibus accession number GSE114853 (human) and GSE114913 (mouse).

For analysis of *Ret*, *Chat*, and *Nos1* transcript expression at the single cell level in the adult mouse ENS, raw data from a recent study in which myenteric neurons were isolated from 6-8 week old Phox2b-CFP reporter mice and sequenced was analyzed[24] (Accession# GSE153202). All raw data from inDrop runs were merged for a total of 3642 neurons from the ileum and 2983 neurons from the colon included in the final analyses, which were performed using the Seurat 4.0.6 package on R. For the human colonic ENS, sequencing data were re-analyzed from a recent study[25] for which raw data were available on the Single Cell Portal for human colon myenteric plexus (droplet-based MIRACL-Seq). The Clustree package was used to identify appropriate clustering resolution. All UMAPs and dotplots were generated with default ggplot graphics software in R.

### Peptide hormone quantitation

For fasting/re-feeding experiments, the final dose of GSK408B or vehicle was administered 30min prior to collection of “fasting” plasma. For fasting plasma collection, submandibular venous blood was collected directly into EDTA-containing tubes with protease inhibitor cocktail (26.1µL Sigmafast protease inhibitor [Sigma-Aldrich], 2.6µL of 2.5µM DPPIV inhibitor [Millipore] and 1.3µL of 100mg/L serine protease inhibitor Pefabloc [Roche]) at a ratio of 11.5µL inhibitor/100µL of blood. For post-prandial plasma, mice were euthanized and blood was collected via cardiac puncture into syringes pre-coated with the cocktail. Samples were spun down at 1000 rcf for 10 minutes at 4˚C, with supernatants collected and frozen on dry ice. To quantify levels of PYY, GLP1 and glucagon, custom Mesoscale U-plex plates were utilized (25µL of plasma per well) and run in duplicate.

### Biogenic amine quantitation

Approximately 100µL of submandibular blood (fasting), 500µL of cardiac blood (post-prandial), or 0.5-1cm of PBS-flushed intestinal tissue was collected into 1.7mL tubes, snap-frozen in liquid nitrogen, and stored at -80°C until analyzed. The Vanderbilt University Neurochemistry Core employed liquid chromatography/mass spectrometry (LC/MS) to determine biogenic amine concentrations. Samples and standards were prepared as previously described[46]. Briefly, tissue samples were homogenized in 0.1M TCA-containing $10^{-2}$ M sodium acetate, $10^{-4}$ M EDTA, and 10.5 % methanol, or 20uL of blood was diluted with 60uL acetonitrile:water (80:20), vortexed, and allowed to sit on ice for 10 minutes. Samples were then spun down and analytes in tissue/blood extract supernatant were quantified following derivatization with benzoyl chloride. LC was carried out on a 2.1 x 100 mm, 1.6µm particle CORTECS Phenyl column (Waters Corporation, Milford, MA, USA) using a Waters Acquity UPLC. Mobile phase A was 0.1% aqueous formic acid, and mobile phase B was acetonitrile with 0.1% formic acid. MS analysis was performed using a Waters Xevo TQ-XS triple quadrupole tandem mass spectrometer. The source temperature was 150°C, and the desolvation temperature was 400°C. Total protein was determined using the BCA Protein Assay Kit (Thermo Scientific, Waltham, MA USA). Results for all analytes are reported in Supplementary Table 3.

**Statistical analyses**

Two-way ANOVAs were used to analyze GITT, FPO, % stool water, fecal pellet mass, proportions of $Ret^{+}$ EEC populations and GLP1/PYY plasma levels with sex and genetic/drug intervention/intestine region/feeding status as independent variables. Two-way ANOVAs were also used to analyze proportion of total neurons with genotype and neuronal subtype/intestinal region as independent variables, and biogenic amine concentrations with treatment or genotype and intestinal region or feeding status as independent variables. Repeated measures ANOVAs were used to assess body mass change over time as well as SITT with genotype/drug treatment as the independent variable. One-way ANOVA was used to compare mean *Ret* mRNA expression in bulk RNA sequencing data from mouse and human datasets independently. $RET^{cKO}$ GITT was analyzed by repeated measures ANOVA with the pre- and post-GITT times the within-subjects factor with genotype and timepoint (2 or 4 weeks) as independent variables. No data points were excluded.

**Supplementary Figure Legends**

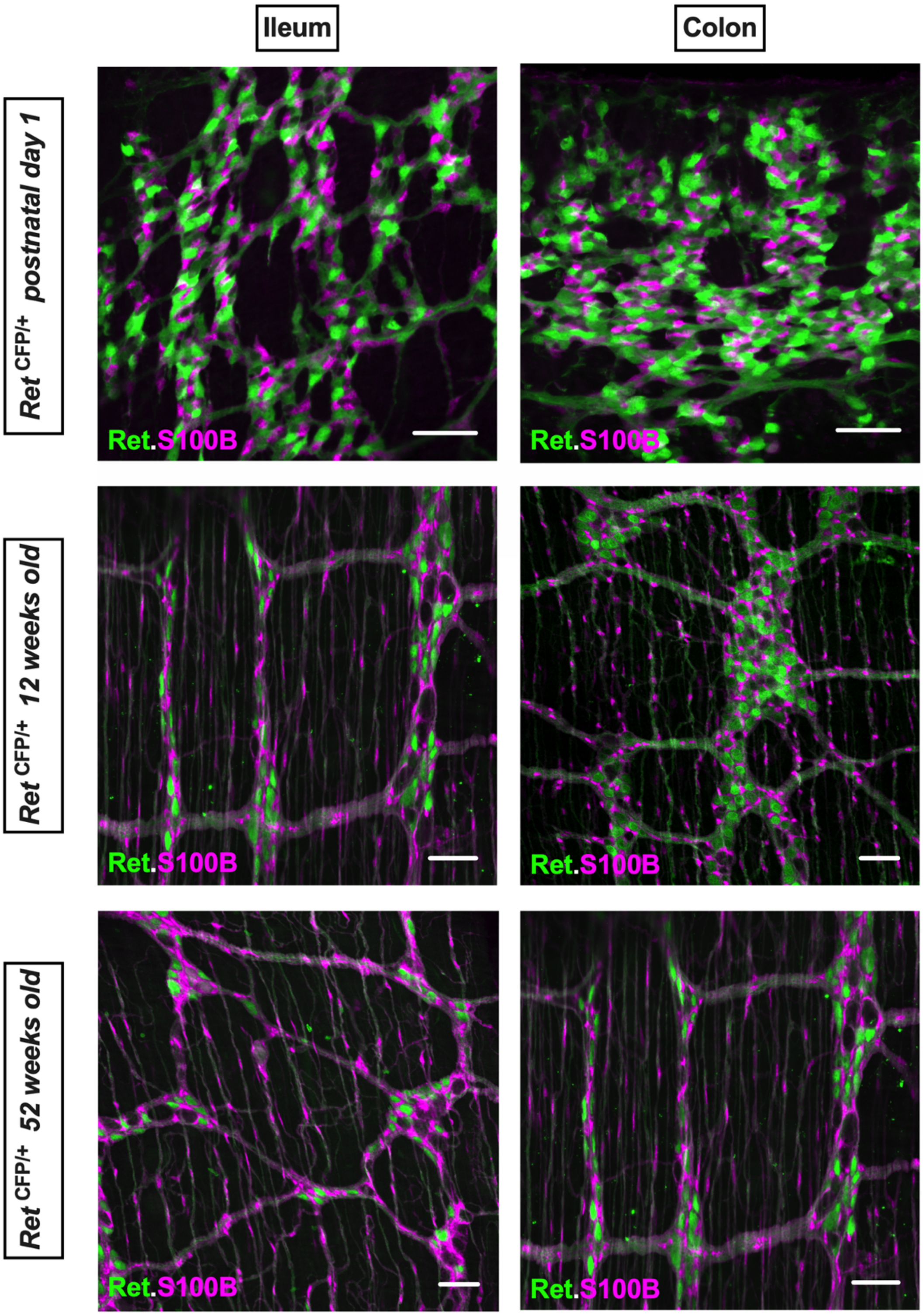


**Supplementary Figure 1. Enteric glia in the postnatal intestine do not express *Ret*.** Representative images of ileum and colon from *Ret*$^{CFP/+}$ mice immunostained as whole mounts and imaged at the level of the myenteric plexus reveal that S100B-immunoreactive enteric glia (magenta) do not express *Ret* (marked by CFP immunoreactivity in green) at 1 day, 12 weeks or 52 weeks of age. *Ret*-expression is not evident in either intraganglionic glia or intramuscular glia outside the plexus. Scale bar = 50μm.

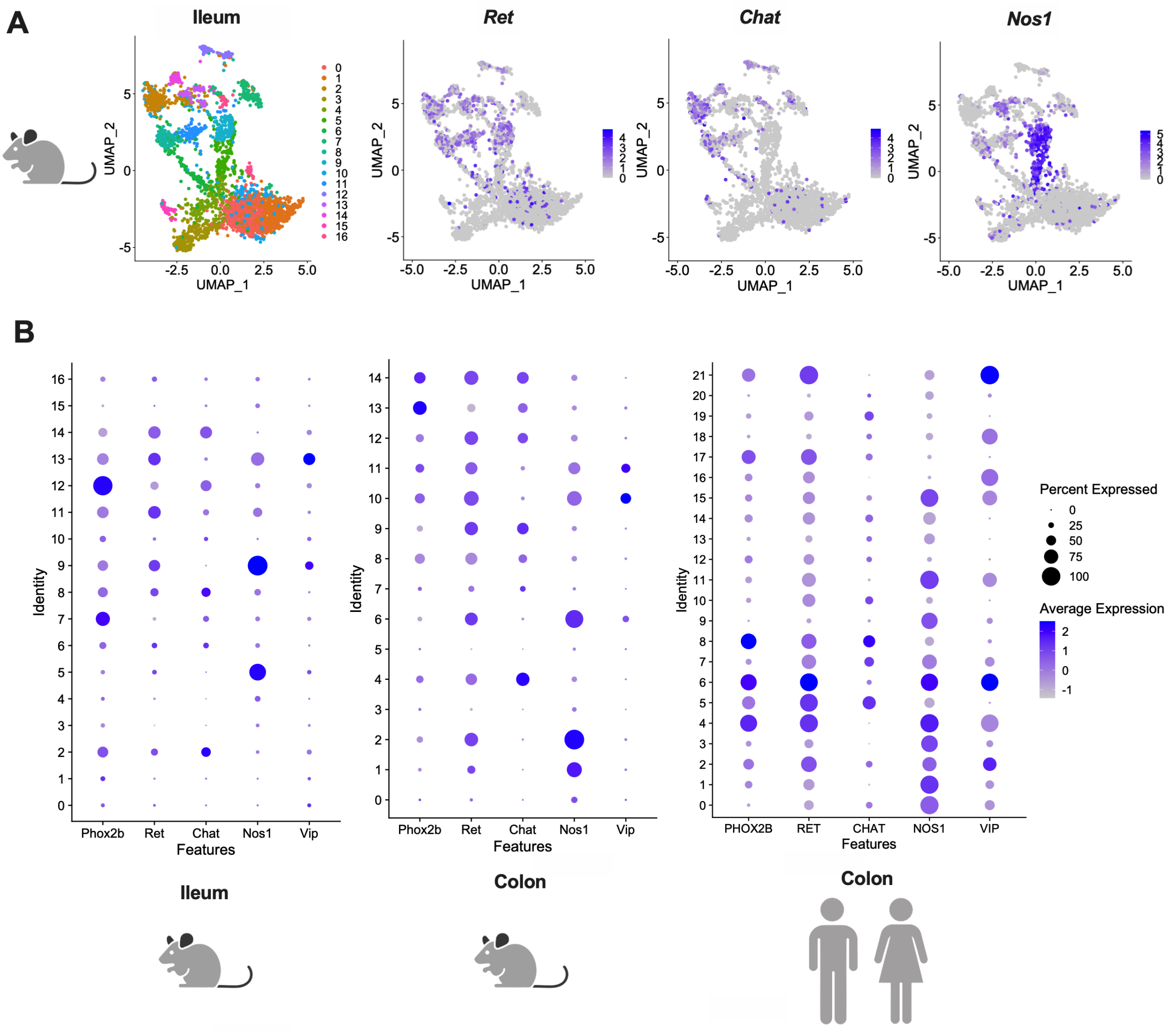


**Supplementary Figure 2. Ret is expressed within subsets of mature enteric neurons in the adult ENS of both mice and humans.**

**(A)** Single cell RNA sequencing of myenteric neurons isolated from adult mouse ileums shows that *Ret* is expressed within a subset of *Chat*$^{+}$ *neurons* and the majority of *Nos1*$^{+}$ neurons (secondary analysis of raw data[24]).

**(B)** Dot-plots of gene expression from adult mouse and human ENS showing distribution and average expression levels of transcripts for *RET*, *CHAT, NOS1* and *VIP* across transcriptionally distinct clusters of neurons. The pan-neuronal gene *PHOX2B* is included for reference (secondary analysis of raw data[24,25]).

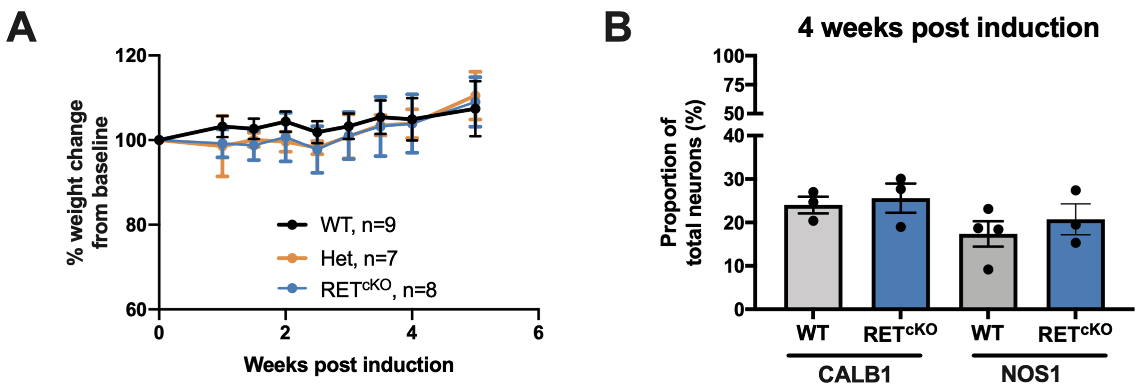


**Supplementary Figure 3. RET depletion in the postnatal gut does not alter body weight or the density of enteric neurons.**

**(A)** Male and female WT, Het and RET$^{cKO}$ mice show no weight loss compared to baseline for over 5 weeks following tamoxifen induction. Each dot represents the group mean and error bars reflect standard deviation.

**(B)** The proportions of CALB1$^{+}$ and NOS1$^{+}$ neurons were unchanged in the ileum and colon, respectively, when measured 4 weeks after tamoxifen administration in RET$^{cKO}$ mice. Graph represents mean ± SEM with individual animals as dots.

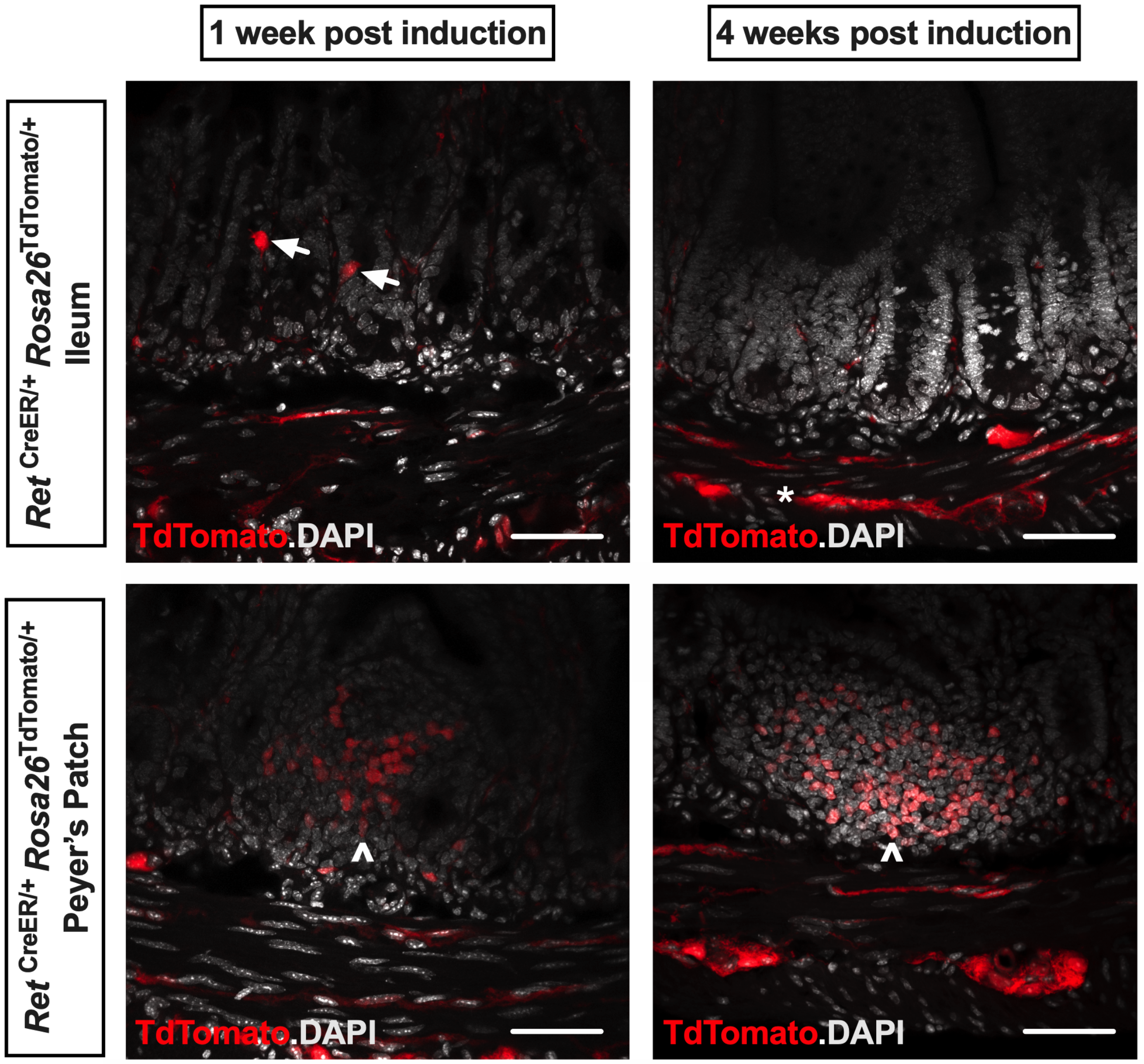


**Supplementary Figure 4. RET is expressed in subsets of neurons, epithelial cells and immune cells in the postnatal gut, which turnover at different rates.**

Representative images of cross-sections of ileum isolated from *Ret*$^{CreER/+}$ *Rosa26*$^{tdTomato/+}$ mice in which *Ret*-expressing cells are labeled with TdTomato reporter upon tamoxifen administration. DAPI marks cell nuclei in white and red marks TdTomato-expressing cells that either expressed *Ret* at the time of tamoxifen exposure or derived from cells expressing *Ret* at that time. At 1-week post-induction, TdTomato expression is evident in neuronal fibers and ganglia (*), isolated epithelial cells (white arrows), and immune cells (arrowheads) in Peyer's patches (PP). At 4-weeks post-induction, neuronal and immune cell TdTomato expression remains robust but epithelial expression is no longer detectable, suggesting that *Ret*-expressing epithelial cells turned over during this period. Scale bar = 50μm

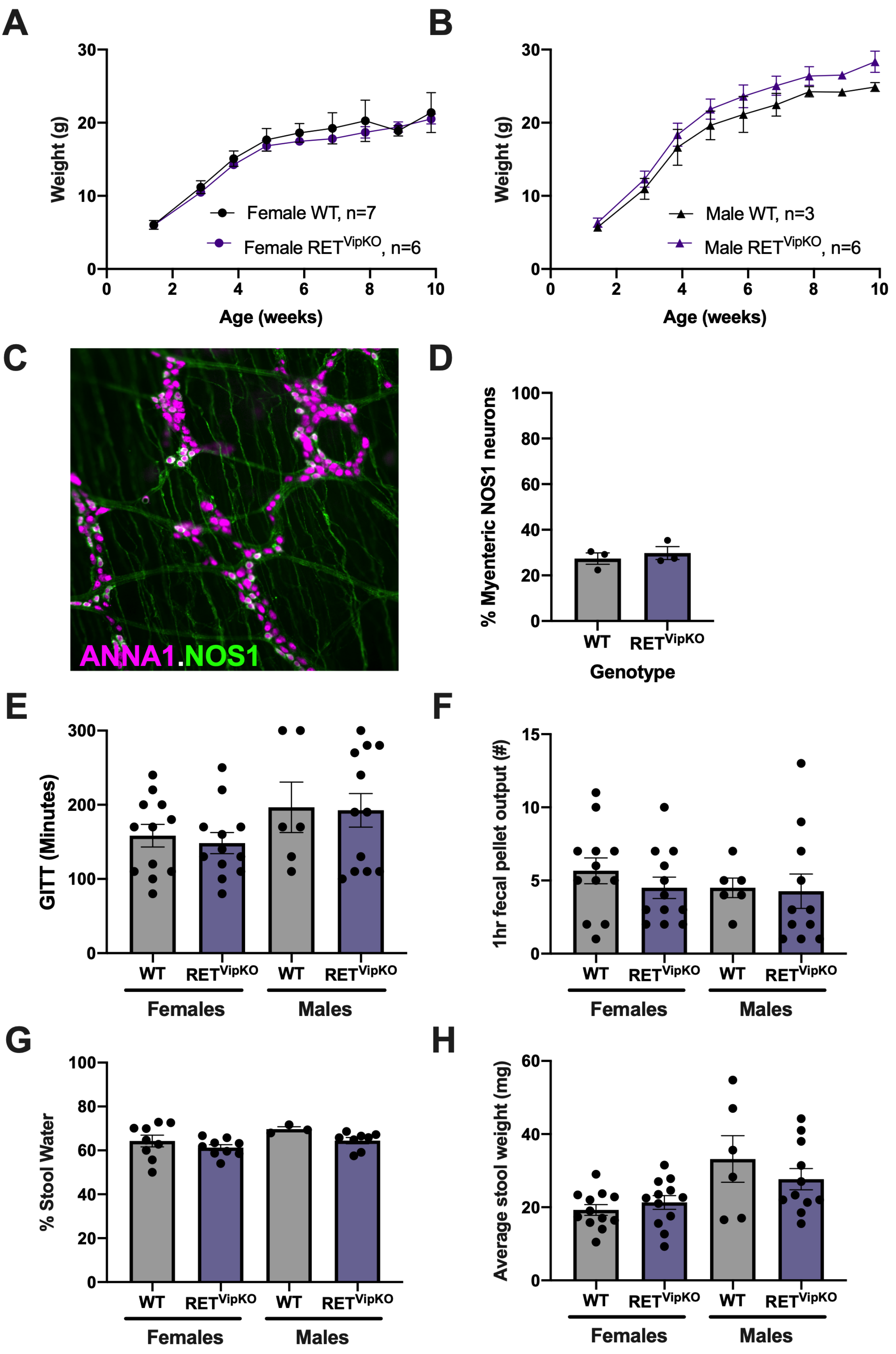
A
Weight (g)
Age (weeks)
Female WT, n=7
Female RETVipKO, n=6
B
Weight (g)
Age (weeks)
Male WT, n=3
Male RETVipKO, n=6
C
ANNA1.NOS1
D
% Myenteric NOS1 neurons
WT
RETVipKO
Genotype
E
GITT (Minutes)
WT
RETVipKO
Females
Males
F
1hr fecal pellet output (#)
WT
RETVipKO
Females
Males
G
% Stool Water
WT
RETVipKO
Females
Males
H
Average stool weight (mg)
WT
RETVipKO
Females
Males

**Supplementary Figure 5. RET$^{VipKO}$ mice exhibit no deficits in body weight, GI motility, fecal composition or density of nitrergic neurons in the myenteric plexus.**

**(A-B)** Body mass in mice lacking RET function in Vip$^{+}$ enteric neurons (*Vip*$^{Cre/+}$*Ret*$^{flx/flx}$; referred to as RET$^{VipKO}$) is no different than that of littermate controls (*Ret*$^{flx/flx}$; referred to as WT) in males or females over time.

**(C-D)** Representative image of ileum from an adult RET$^{VipKO}$ mouse immunostained as whole mounts and imaged at the level of the myenteric plexus shows that NOS1-immunoreactive neurons (green) appear grossly intact and represent a similar proportion of all enteric neurons (labeled by ANNA-1) as in WT controls.

**(E-F)** Gastrointestinal transit time (GITT) and 1-hour FPO are no different between RET$^{VipKO}$ mice and controls, among males or females.

**(G-H)** The water content (%) and average mass of spontaneously expelled fecal pellets were no different between RET$^{VipKO}$ mice and controls, among males or females.

**A** and **B** are presented as mean ± SD, with each dot representing a group mean. **D-H** are presented as mean ± SEM with individual animals as dots.

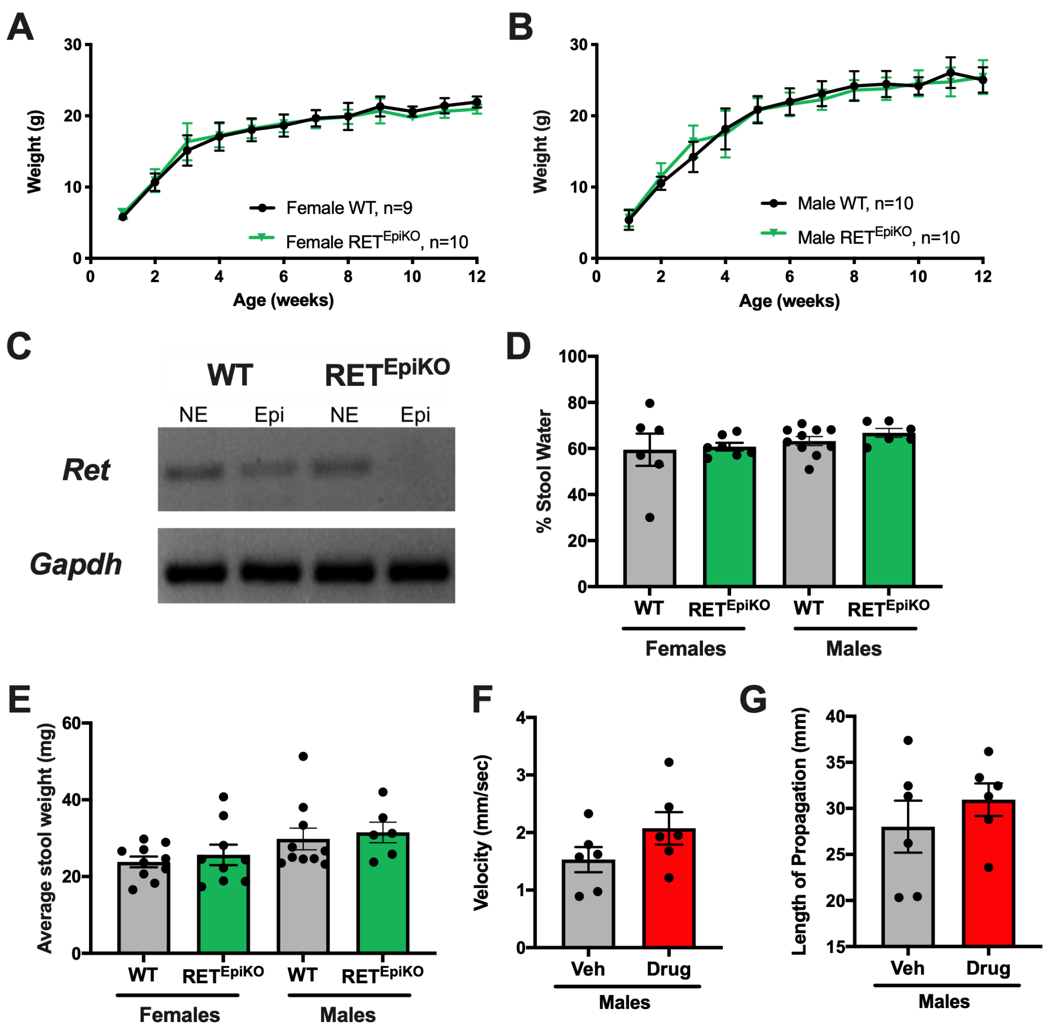


**Supplementary Figure 6. Disruption of epithelial RET signaling does not alter body weight or measures of colonic function.**

**(A-B)** Body mass in mice lacking RET function in the intestinal epithelium (*Vil1*$^{Cre}$ *Ret*$^{flx/flx}$; referred to as RET$^{EpiKO}$) is no different than that of littermate controls (*Ret*$^{flx/flx}$; referred to as WT) in males or females over time.

**(C)** Representative image of RT-PCR shows that RET$^{EpiKO}$ mice exhibit selective loss of *Ret* transcript expression in the epithelial layer of the small intestine (Epi), while expression in the non-epithelial layers (NE) is similar to that in WT controls.

**(D-E)** The water content (%) and average mass of spontaneously expelled fecal pellets were no different between RET$^{EpiKO}$ mice and controls, among males or females.

**(F-G)** Related to Figure 3E, colonic motor behaviors were observed by *ex vivo* video imaging of colons isolated from male mice treated with a gut-restricted RET kinase inhibitor (Drug) or vehicle (Veh) for 3.5 days. The velocity and length of propagation of colonic migrating motor contractions (CMMCs) were not altered by RET kinase inhibition.

**A** and **B** are presented as mean ± SD, with each dot representing a group mean. **D-G** are presented as mean ± SEM with individual animals as dots.

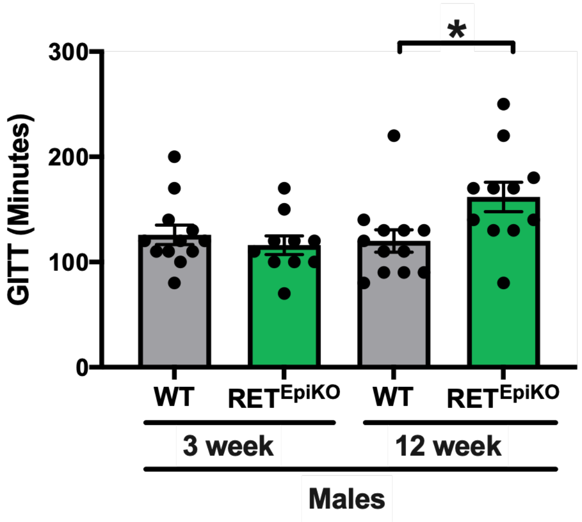


**Supplementary Figure 7. Epithelial RET regulation of GI motility is age-dependent in male mice.**

At 3 weeks of age, just after weaning and before puberty, male *Ret*$^{EpiKO}$ mice have the same GI transit times (GITT) as control littermates. At 12 weeks of age, well after puberty, GITT is 30% slower in *Ret*$^{EpiKO}$ males compared to controls. Data presented as mean ± SEM with individual animals as dots. * indicates $p<0.05$ by two-way ANOVA.

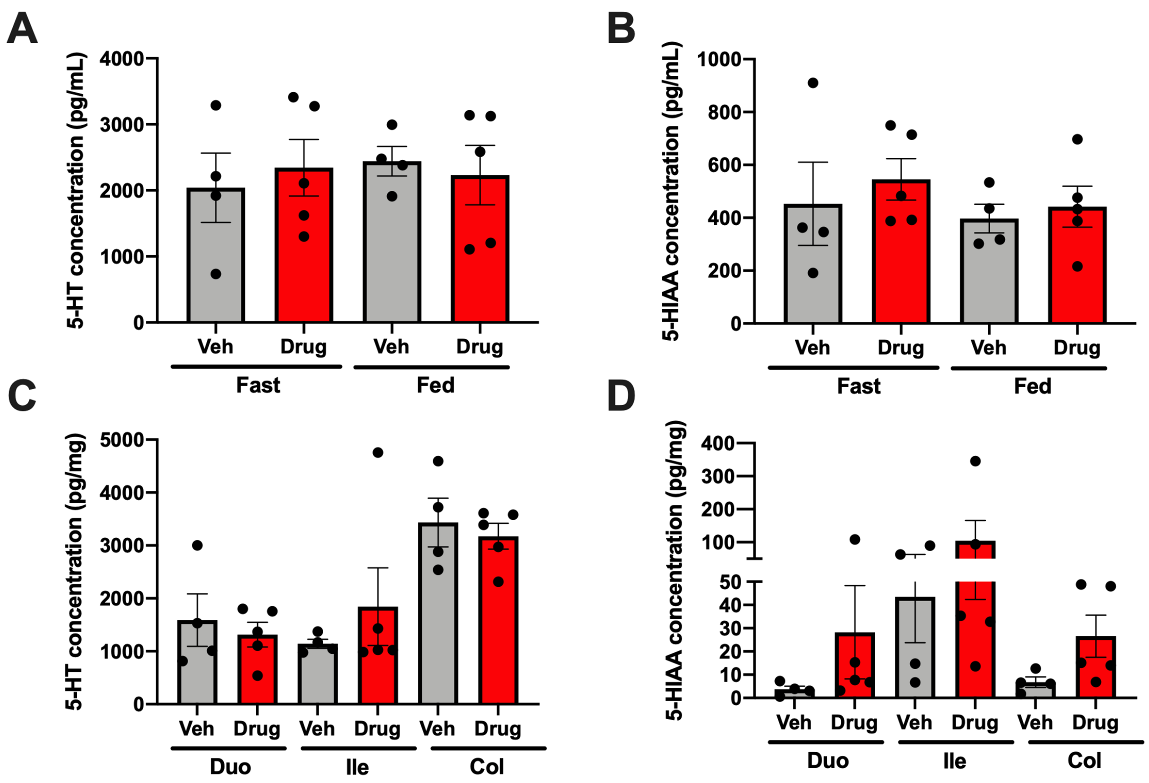


**Supplementary Figure 8. RET kinase inhibition does not alter blood or tissue levels of 5HT or 5-HIAA.**

(**A-B**) Male mice treated with a RET kinase inhibitor do not exhibit altered levels of 5-HT (A) or 5-HIAA (B) in whole blood in either fasted or fed states. No differences are seen between mice treated with the Ret kinase inhibitor (Drug, red) and those treated with vehicle (Veh, grey).

(**C-D**) RET kinase inhibition in male mice does not alter the levels of 5-HT (C) or its major metabolite 5-HIAA (D) in the duodenum (Duo), ileum (Ile) or colon (Col).

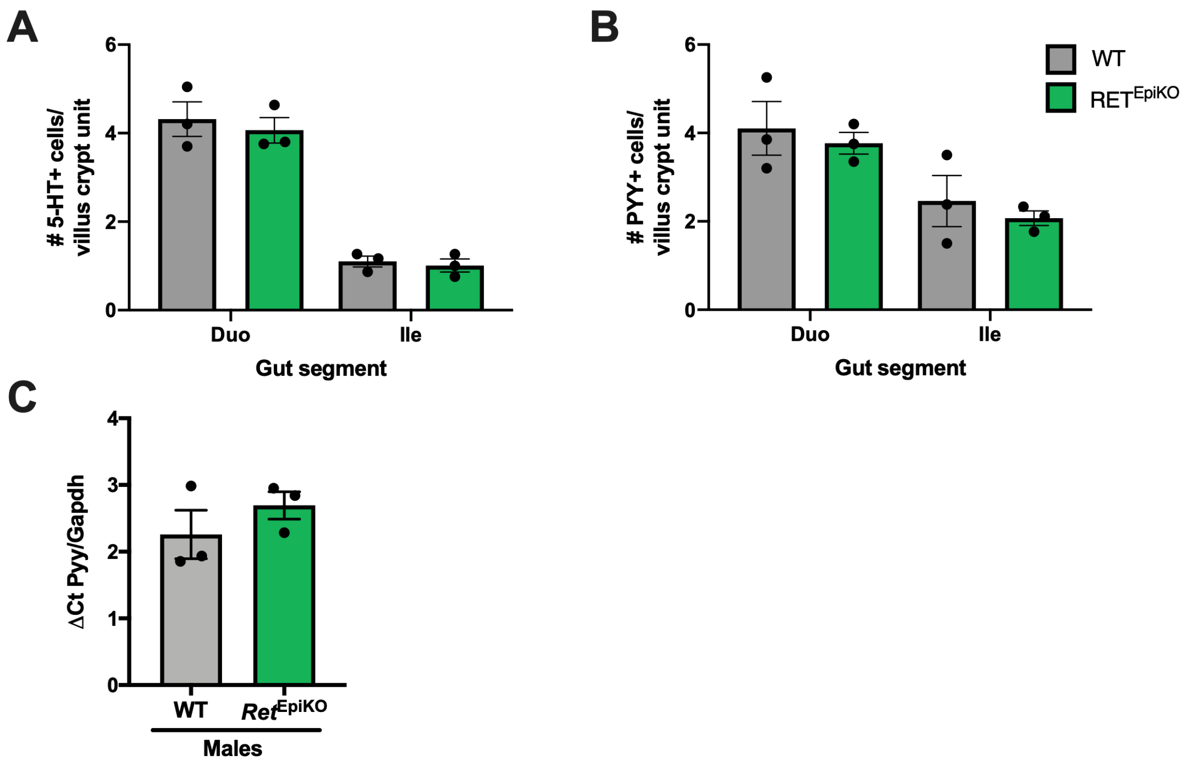


**Supplementary Figure 9. Genetic disruption of epithelial RET does not alter the density of enterochromaffin or L-cells in the small intestine.**

**(A-B)** RET$^{EpiKO}$ mice have similar numbers of 5-HT-immunoreactive cells (A) and PYY-immunoreactive cells (B) per villus-crypt unit in the duodenum (Duo) and Ileum (Ile) as WT littermates.

**(C)** *Pyy* transcript levels in the small intestinal epithelium are no different in WT and RET$^{EpiKO}$ mice.

All graphs are presented as mean ± SEM with values from individual animals as dots.

**Supplementary Table 1. Antibodies used for immunohistochemistry experiments**

| Reagent | Source | Identifier | Concentration |
|---|---|---|---|
| Rabbit anti-GFP | Invitrogen | A11122 | 1:1000 |
| Chicken anti-GFP | Aves | GFP-1020 | 1:1000 |
| Human anti-ANNA | Mayo Clinic | Gift from V. Lennon | 1:40,000 |
| Rabbit anti-NOS1 | Immunostar | 24287 | 1:1500 |
| Rabbit anti-CALB1 | Swant | CB38 | 1:400 |
| Rabbit anti-S100B | DAKO | Z0311 | 1:500 |
| Rabbit anti-DsRed | Takara | 632496 | 1:500 |
| Rabbit anti-CHGA | Abcam | ab15160 | 1:1000 |
| Rabbit anti-5HT | Immunostar | 20080 | 1:1000 |
| Rabbit anti-PYY | Bioss USA | bs-2265R | 1:500 |

**Supplementary Table 2. Sequences of primers used for PCR and qRT-PCR experiments**

| | **5' to 3' sequence** | |
|---|---|---|
| **Gene** | **Forward Primer** | **Reverse Primer** |
| Ret | CAGGATGGGCCACTTCTTCT | CACATAGGCAGGCCCAATCT |
| Rpl19 | ACCTGGATGAGAAGGATGAG | ACCTTCAGGTACAGGCTGTG |
| Pyy | TTCACAGACGACAGCGACAG | CACCACTGGTCCAAACCTTCT |
| Gapdh | ATGTGTCCGTCGTGGATCTGA | GCTGTTGAAGTCGCAGGAGACA |

**Supplementary Table 3. Tissue levels of biogenic amines in male mice exposed to RET kinase inhibitor are no different than those in mice exposed to vehicle.**
N = 4-5 adult male mice per condition.

| Group | Intestinal segment | Dopamine | | 5HT | | 5HIAA | | Glutamate | | NE | |
|---|---|---|---|---|---|---|---|---|---|---|---|
| | | Mean | SEM | Mean | SEM | Mean | SEM | Mean | SEM | Mean | SEM |
| Vehicle | Duo | 47.06 | 36.36 | 1590.3 | 494.54 | 3.74 | 1.38 | 7117.55 | 924.77 | 61.65 | 18.49 |
| Drug | Duo | 9.02 | 2.94 | 1315.44 | 232.48 | 28.24 | 20.07 | 6245.81 | 1542.8 | 56.11 | 7.37 |
| Vehicle | Ile | 13.75 | 5.77 | 1141.68 | 86.34 | 43.5 | 19.75 | 7089.11 | 321.19 | 97.76 | 13.33 |
| Drug | Ile | 7.77 | 1.53 | 1115.95 | 104.97 | 104.19 | 61.84 | 5493.34 | 993.49 | 92.11 | 11.3 |
| Vehicle | Col | 8.48 | 4.23 | 3434.4 | 459.62 | 6.8 | 2.25 | 3517.77 | 339.75 | 81.47 | 21.63 |
| Drug | Col | 16.07 | 12.26 | 3174.1 | 243.36 | 26.57 | 9.04 | 5811.58 | 2319.14 | 78.43 | 10.45 |
